# Axon length-dependent synapse loss is mediated by neuronal cytokine-induced glial phagocytosis

**DOI:** 10.1101/2024.06.09.598122

**Authors:** Federico Tenedini, Chang Yin, Jessica Huang, Neena Dhiman, Peter Soba, Jay Z. Parrish

## Abstract

Many neurodegenerative disorders (NDDs) preferentially affect neurons with long or complex axonal arbors, but our understanding of this specific vulnerability is limited. Using *Drosophila* larval class IV dendrite arborization (C4da) neurons, we found that neuronal activation of the integrated stress response (ISR) induces axon length-dependent degeneration (LDD). We identified the Interleukin-6 homologue unpaired 3 (upd3) as both necessary and sufficient for LDD in C4da neurons. Upd3 recruits glial cells to phagocytose presynapses preferentially on neurons with long axons, revealing an intrinsic axon length-dependent vulnerability to glia-mediated presynapse removal. Finally, we found that axon length-dependent presynapse loss in fly models of human NDDs utilized this pathway. Altogether, our studies identify inflammatory cytokine signaling from neurons to glia as a key determinant in axon length-dependent vulnerability.

**One-Sentence Summary:** Sensory neurons exhibit intrinsic length-dependent vulnerability to presynapse removal driven by cytokine activation of glia.

## Introduction

Neurodegenerative disorders (NDDs) affect approximately 15% of the world’s population (*1*), creating a major health burden for aging societies in all industrialized countries. A majority of NDDs are idiopathic with age as the biggest risk factor, but multiple different genetic loci are linked to early onset degeneration as well, confounding treatment (*2*). While the primarily affected neuronal subsets differ, common pathological features of many NDDs provide opportunities for therapeutic intervention (*3*). One particularly salient feature of NDDs is the anatomical complexity of affected cells: neurons with extensive projections are more prone to degeneration, and the severity of degenerative phenotypes often correlates with axon length. This length-dependency is best illustrated in neuropathies including Charcot-Marie-Tooth (CMT) disease, hereditary spastic paraplegia (HSP) and amyotrophic lateral sclerosis (ALS), which show length-dependent degeneration of peripheral axons typically starting in distally innervated regions of the body (*4–8*). Similarly, peripheral neuropathy in some forms of spinocerebellar ataxia (SCA) involves length-dependent axonopathy (*9–11*). However, central nervous system neuron populations with large axon length and/or arbor complexity are likewise primary targets of degeneration: cholinergic cells in the basal forebrain are thought to be affected early in Alzheimer’s disease (AD) and feature extensive axonal arborization estimated to reach 100m per cell (*11–13*); upper motor neurons innervating distal spinal cord targets are commonly affected in ALS (*5*); dopaminergic neurons of the substantia nigra pars compacta (SNc) have the largest arbors and highest number of synapses of all dopaminergic neurons (*5*), and are the most strongly affected subset and primary cause of motor defects in Parkinson’s disease (PD) (*14*, *15*). They are also affected in the most common autosomal dominant form of SCA caused by polyglutamine repeat expansion of ATXN3 (SCA3) (*16*, *17*). Moreover, mouse models link selective vulnerability of SNc dopaminergic neurons directly to their axon arborization size (*18*).

During early NDD pathogenesis, synapse loss is thought to precede axon degeneration and subsequent neuronal cell death as observed, for example, in ALS (*19*), PD(*20*, *21*), and AD (*22*). However, remarkably little is known about the mechanisms required for maintenance of these large neurons and axon length-dependent vulnerability, synapse loss, and degeneration. In this study, we developed an experimental system for systematic dissection of axon length-dependent degeneration using *Drosophila* nociceptive C4da neurons, which are optically and genetically accessible and morphologically well-defined.

They feature stereotyped arbors distinguished only by increasing axon length based on their position along the anterior to posterior body axis. Their synaptic targets and precise number of synapses are defined, facilitating quantitative analysis of connectivity. Furthermore, these neurons mediate stereotyped responses to noxious stimuli, providing robust behavioral outputs to probe neuronal network function. Using this system, we identify a signaling axis initiated by activation of the integrated stress response (ISR) in neurons which drives production of inflammatory cytokines, glial recruitment, and ultimately axon length-dependent synapse loss and neuronal dysfunction.

## Results

### C4da neurons show axon length-dependent and -independent degeneration

We aimed to investigate whether specific cellular features, in particular axon length, give rise to vulnerability under neurodegenerative conditions in a single class of neurons. The *Drosophila* larval body is composed of 12 segments, each of which has a similar complement of somatosensory neurons (SSNs). Within this body plan, two major factors impact overall neurite length of SSNs. First, different SSN cell types within a given segment exhibit a ∼10-fold range in peripheral arbor length (*23*). Second, SSN axons innervate the neuropil during embryogenesis and SSNs located in posterior segments must extend their axons further to maintain connectivity during larval growth (Fig. 1A). We focused our studies on the SSNs with the most expansive arbors, the nociceptive C4da neurons. Despite their differences in axon length, C4da neurons in each abdominal segment have comparably sized terminal axonal domains (Fig. 1B) that form equivalent numbers of synapses with identified postsynaptic partners and total numbers of presynapses overall (*24*), providing a sensitive and highly quantitative system for studying axon length-dependent degeneration (LDD). We previously identified *pathetic* (*path*), which encodes a putative amino acid transporter, as a factor that imposes an upper limit on overall neurite length in larval SSNs (*25*, *26*)and therefore reasoned that *path* mutants may provide an entry point to understanding mechanisms of LDD.

**Fig. 1.**
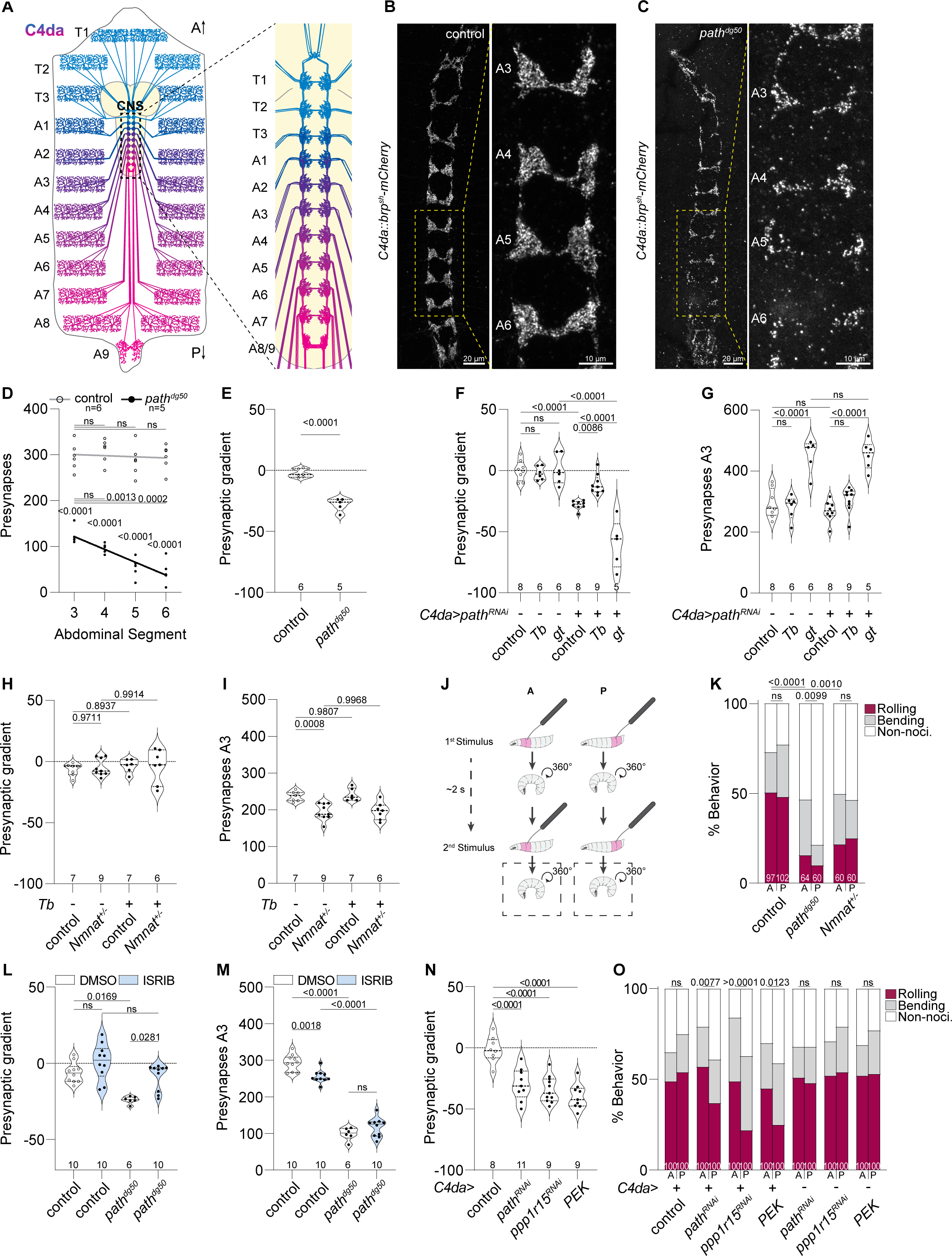
Axon length-dependent degeneration of C4da neuron synapses. **(A)** Schematic depicting the position of C4da nociceptive sensory neurons in a larval filet, shaded according to increasing length of axon projections to the larval CNS. Note that C4da neurons innervating the most posterior segments have the longest axons. Larval segments are indicated (T, thoracic; A, abdominal). **(B-G)** *path* mutation elicits axon length-dependent effects on C4da neuron presynapses. Maximum intensity projections depict the distribution of C4da neuron presynapses labeled by *ppk::brp^sh^-mCherry* in the VNC of control **(B)** and *path^dg50^* mutant **(C)** third instar larvae. ROIs depict presynaptic domains from segments A3-A6, which are used for quantitative analysis throughout this study. **(D)** *path* mutation induces a gradient of presynapses. Plot depicts quantification of presynaptic spots from control and *path^dg50^* mutant larvae. *path* mutation induces a significant reduction in presynapses that is more severe in posterior segments. Lines depict linear regression for control (gray) and *path^dg50^* (black) datasets. P values, ANOVA with post-hoc Šídák’s multiple comparisons test. **(E)** Plot depicts gradient of presynapses from segment A3 to A6 in control and *path^dg50^* mutant larvae. *path* mutants exhibit a negative presynaptic gradient, reflecting the decreasing number of presynapses from A3 to A6. P < 0.0001, unpaired t-test. **(F-G)** Severity of presynapse defects induced by *path* inactivation is dependent on axon length. Violin plots depict presynapse gradient measurements **(F)** and segment A3 presynapse numbers **(G)** from control larvae or larvae with C4da-specific *path* knockdown in combination with mutations in *Tb* to shorten or *gt* to lengthen larvae. *Tb* suppresses and *gt* enhances presynaptic gradient defects. P values, ANOVA with post-hoc Šídák’s multiple comparisons test. **(H-I)** *Nmnat^+/-^* mutation induces axon length-independent degeneration. Violin plots depict presynapse gradient measurements **(H)** and segment A3 presynapse numbers **(I)** from control or *Nmnat^+/-^* heterozygotes in combination with mutation in *Tb* to shorten or *gt* to lengthen larvae. P values, ANOVA with post-hoc Šídák’s multiple comparisons test. **(J)** Mechanonociception behavior paradigm. Two successive mechanical stimuli are delivered to anterior (A) or posterior (P) segments, and larval responses to the second stimulus are recorded. **(K)** LDD triggers more severe functional defects in posterior segments. Histogram depicts mechanonociceptive behavior responses of larvae of the indicated gentoypes. *path^dg50^* but not *Nmnat^+/-^* results in decreased behavior in posterior segments compared to anterior. P-values, Chi-square test. **(L-M)** ISR contributes to *path* mutant LDD phenotypes. Violin plots depict presynapse gradient measurements **(L)** and segment A3 presynapse numbers, P values, Kruskal-Wallis test with post-hoc Dunn’s correction for multiple comparisons, **(M)** from control and *path* mutant larvae fed with DMSO or the ISR inhibitor ISRIB. ISRIB feeding ameliorates *path* mutant LDD but not LID defects. P values, ANOVA with post-hoc Šídák’s multiple comparisons test. **(N)** Activating ISR in C4da neurons triggers LDD. Violin plot depicts presynapse gradient measurements from control larvae, larvae with C4da-specific *path^RNAi^*, or larvae with C4da-specific ISR activation (via *ppp1r15 ^RNAi^* or *PEK* overexpression). A3 presynapse numbers are plotted in fig. S3F. P values, ANOVA with post-hoc Dunnett’s multiple comparisons test. **(O)** ISR activation in C4da neurons triggers axon length-dependent deficits in mechanonociception. Plot depicts behavioral responses of larvae of the indicated genotypes to anterior and posterior noxious mechanical stimuli. P values, Chi-square test. Sample numbers are shown for each specimen and detailed experimental genotypes are provided in table S4.

To assay for length-dependent effects of *path* mutation on C4da axons, we selectively expressed the presynaptic marker *brp^sh^::mCherry* in C4da neurons and monitored presynapse numbers in different abdominal segments of the ventral nerve cord (VNC). Consistent with prior reports (*24*), we found that C4da axons in control larvae exhibited comparable numbers of presynapses in each abdominal segment (segmental means: A3: 296.1 ± 27.2, A4: 309 ± 23.6, A5: 288 ± 30.5, A6 294.8 ± 25.1) (Fig. 1B, D). In contrast, larvae homozygous for a null mutation in *path* (*path^dg50^*) exhibited a significant reduction in presynapses that was both graded along the anterior-posterior (AP) axis and progressive (Fig. 1, C and D, and fig. S1A). *path* mutants exhibited significantly fewer presynapses in each segment when compared to controls, while additionally displaying significantly fewer presynapses in the posterior compared to anterior segments (Fig. 1, B to D). We quantified this axon length-dependent gradient of presynapse loss by measuring the slope of the linear regression fit to the number of presynapses in the segmental domains (Fig. 1E). Lastly, although *path* mutant and control larvae exhibited comparable numbers of C4da presynapses early during larval development, *path* mutant larvae showed a progressive reduction in presynapses after the 48 h timepoint (fig. S1A). This indicates that *path* mutation not only restricts presynapse addition but leads to an active decline of presynaptic structures during larval growth.

*Path* functions cell autonomously in SSNs to regulate neuron growth but additionally, functions in glial cells to promote brain sparing during nutrient restriction (*27*). We therefore used neuron-specific RNAi to assess cell-autonomous *path* function in axon maintenance. We found that *path^RNAi^* in C4da neurons significantly reduced presynapse numbers in neurons innervating posterior but not anterior segments, resulting in a presynaptic gradient comparable in extent to *path^dg50^* mutant larvae (Fig. 1, F and G). To assess the axon length-dependency of *path* mutant presynapse phenotypes, we genetically manipulated larval size and hence axon length by shortening or lengthening the animals. We found that mutations in *Tubby* (*Tb^1^*) and *giant* (*gt^1^*) yielded a 19% reduction and 85% increase in larval body length overall and exhibited graded effects on axon length (fig. S1, B to D). While *Tb^1^* had no effect on C4da presynapse numbers on its own, *gt^1^* larvae displayed continuous presynapse addition during an extended larval period and hence an overall increase in presynapse numbers (Fig. 1G). Remarkably, *Tb^1^* suppressed and *gt^1^* enhanced the presynaptic gradient triggered by *path* knockdown (Fig. 1F), further underscoring the relationship between axon length and presynapse deficits in neurons lacking *path* function. Altogether, these results indicate that *path* is required cell-autonomously in C4da neurons to maintain presynapse numbers, particularly in neurons with the longest axons.

### LDD is distinct from mechanisms of axon length-independent degeneration

Axon degeneration in many NDDs involves mechanisms related to Wallerian degeneration (WD) (*28–32*). To assess the potential contributions of WD to LDD, we utilized mutations in the conserved Nicotinamide mononucleotide adenylyltransferase *Nmnat*. Homozygous *Nmnat* mutations induce early larval lethality, but heterozygous *Nmnat* mutations trigger dendrite and axon degeneration in larval C4da neurons (*33*, *34*). Similarly, larvae heterozygous for a *Nmnat* null mutation (*Nnmat^+/-^*) exhibited a significant reduction in C4da presynapse numbers (Fig. 1H). However, the *Nnmat^+/-^* presynapse loss phenotype was distinct from that observed in *path* mutants in two important ways. First, *Nnmat^+/-^* larvae exhibited a comparable extent of presynapse loss in anterior and posterior segments and hence exhibited no presynapse gradient (Fig. 1I and fig. S2A). Second, axon shortening induced by *Tb^1^* had no effect on the *Nnmat^+/-^* induced presynapse loss (Fig. 1, H and I). Taken together, these results indicate that presynapse loss induced by *Nmnat* mutation is independent of axon length, a phenomenon we refer to as axon length-independent degeneration (LID) (fig. S2B).

Next, to assay functional consequences of LDD and LID, we measured larval nocifensive responses (rolling/bending) to mechanical nociceptive cues delivered to either anterior (A2/A3) or posterior (A6/A7) segments in control, *path^dg50^* and *Nmnat^+/-^* larvae. For these studies we monitored responses to two successive stimuli (Fig. 1J), a paradigm that generates robust and reproducible nocifensive responses (*35*, *36*). Whereas anterior and posterior stimulation yielded comparable behavioral responses in control larvae, *path^dg50^* and *Nmnat^+/-^* mutant larvae exhibited reduced mechanonociceptive responses to stimuli at both sites, consistent with the overall reduction in C4da presynapses caused by each mutation (Fig. 1K). However, *path^dg50^* but not *Nmnat^+/-^* larvae displayed significantly less nocifensive behavior to stimulation in posterior segments. Hence, the graded C4da presynapse loss observed in LDD is accompanied by graded functional deficits in nociception.

### Activation of the integrated stress response induces LDD

*Path* inactivation triggers cellular responses associated with nutrient deprivation including mitochondrial fusion and reduced translational capacity (*25*). Phosphorylation of the alpha subunit of eukaryotic initiation factor 2 (P-eIF2α) is a nexus for translational control by the integrated stress response (ISR), an adaptive pathway that promotes cell survival in response to a variety of stressors and is deregulated in NDDs affecting large neurons (*37–39*). We therefore examined connections between *path* mutation, the ISR and LDD. First, we compared P-eIF2α levels in control and *path* mutant larvae and found that *path* inactivation increased P-eIF2α levels in C4da neurons without affecting eif2α levels overall (fig. S3, A and B). Next, we assayed for contributions of ISR activation to *path* mutant LDD phenotypes using ISRIB, a compound which antagonizes translational inhibition triggered by eIF2α phosphorylation (*40*, *41*). On its own ISRIB feeding had no major effect on presynapse numbers (Fig. 1M and fig. S3C), but ISRIB feeding in *path* mutants eliminated the presynaptic gradient (Fig. 1L), doing so by increasing posterior (A6) but not anterior (A3) presynapse numbers in a dose-dependent manner (Fig. 1M and fig. S3C). ISRIB feeding likewise eliminated the posterior hyposensitivity to noxious mechanical stimuli triggered by *path^dg50^* mutation without affecting controls or anterior responses (fig. S3D). In contrast, constitutive ISR activation in C4da neurons via overexpression of the eif2α kinase *PEK* or inactivation of the PEK antagonist *ppp1R15* yielded LDD phenotypes comparable to those seen with *path* inactivation and additionally induced LID (Fig. 1, N and O; fig. S3, E and F). These results indicate that ISR activation in C4da neurons is necessary for LDD in *path* mutants, and that activation of ISR is sufficient to trigger LDD.

### Neuronal expression of the inflammatory cytokine upd3 triggers LDD

To identify putative mediators of LDD, we used RNA-seq to examine transcriptional changes induced by *path* inactivation or ISR activation in C4da neurons (Fig. 2A and fig. S4, A to D). We identified a set of 109 differentially expressed genes (DEGs) that were significantly altered in response to each of these treatments (Fig. 2B and table S1). This set of DEGs was enriched for genes implicated in NDDs, including genes involved in cellular responses to oxidative stress (fig. S4E), hence we hypothesized that an imbalance in neuronal redox potential contributes to the degenerative phenotypes. To test this possibility, we utilized a genetically encoded fluorescent sensor to monitor the Glutathione (GSH) redox status in C4da neurons and found that the cytosolic and mitochondrial GSH redox potential was significantly reduced in axonal termini of *path^dg50^* mutant larvae (fig. S5, A to D). Next, we experimentally manipulated the redox potential in C4da neurons and assayed effects on LDD phenotypes. C4da-specific knockdown of *Glutathione S transferase S1 (GstS1),* which was downregulated in conditions that induced LDD (Fig. 2B), induced LDD phenotypes (fig. S5E). In contrast, ectopic expression of *Catalase* (*Cat*), which encodes an antioxidant enzyme that catalyzes hydrogen peroxide breakdown, suppressed LDD induced by C4da-specific *path* inactivation (fig. S5F). Taken together, these results demonstrate that oxidative stress is linked to the axon length-dependent vulnerability of presynapses in C4da neurons.

**Fig. 2.**
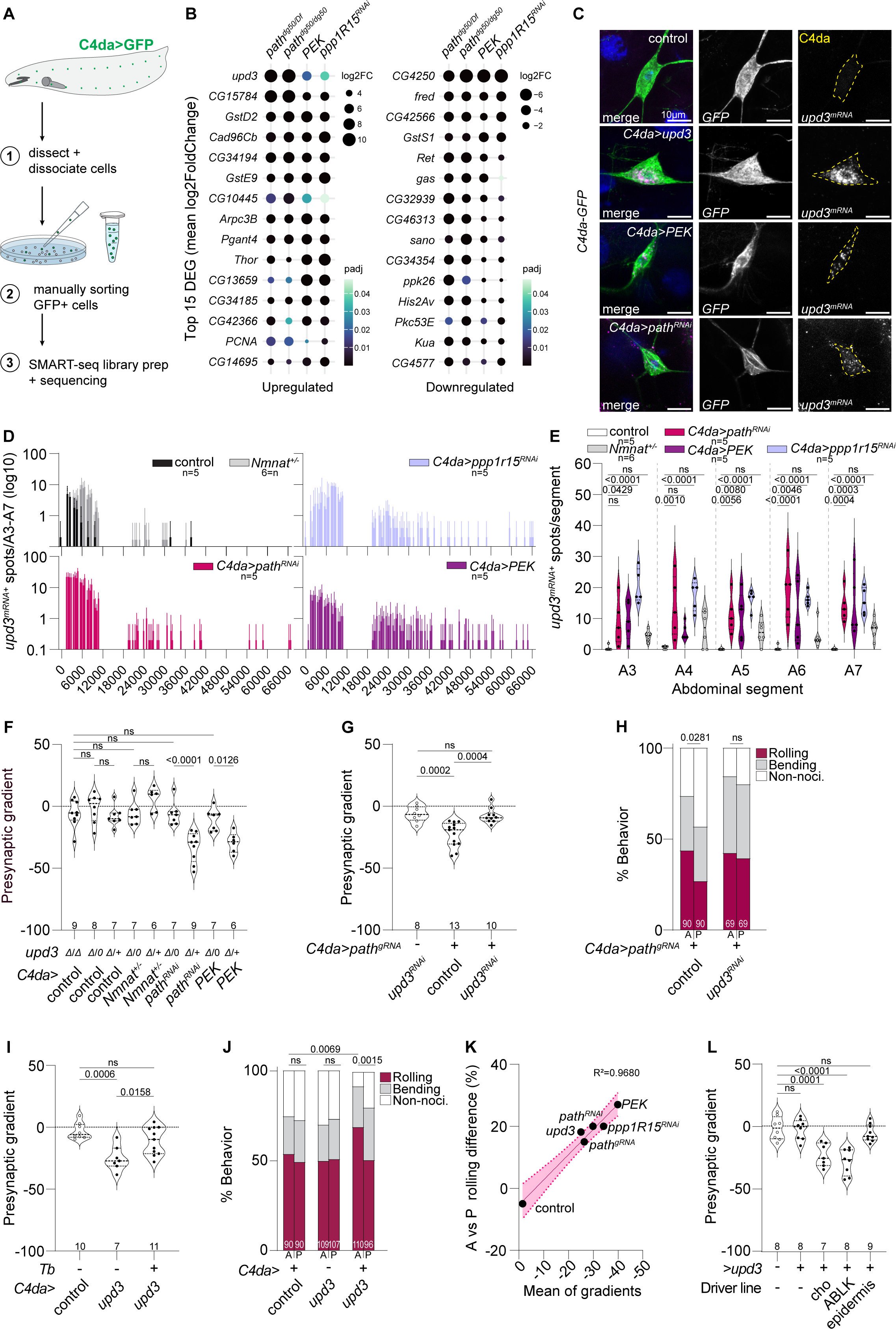
Neuronal expression of the inflammatory cytokine *upd3* drives LDD. **(A-B)** RNA-seq analysis of C4da neurons undergoing LDD. **(A)** Workflow for SMART-seq analysis of 10-cell pools of dissociated, manually sorted C4da neurons. **(B)** Differential expression analysis comparing C4da neuron transcriptomes of wild-type controls and larvae undergoing LDD (two allelic combinations of *path*; *PEK* overexpression in C4da neurons; *ppp1R15* ^RNAi^ in C4da neurons) revealed a set of 109 DEGs deregulated across all LDD conditions. Dot plots depict the top 15 DEGs that were upregulated (left) and downregulated (right) across treatment groups. Dot diameter indicates the mean log_2_FC value across samples and dot shading indicates the adjusted p-value from differential expression analysis. **(C)** HCR-FISH analysis of *upd3* expression in C4da neurons. Representative maximum intensity projections depict staining with Hoechst to label nuclei, GFP antibodies to label C4da neurons (*ppk-8xmCD8-tdGFP*), and *upd3* HCR-FISH fluorescence signal in the following larval fillet preparations: wild-type control (top), C4da overexpression of *upd3* (*C4da>upd3*), C4da overexpression of *PEK* (*C4da>PEK*), and C4da RNAi of *path* (*C4da>path^RNAi^*). **(D-E)** *upd3* mRNA expression is induced in C4da neurons undergoing LDD but not LID. **(D)** Histograms depict the number and intensity of *upd3* HCR-FISH puncta in C4da neuron soma of the indicated genotypes. **(E)** Violin plots depict the number of *upd3* puncta with intensity > 8000 AU in each abdominal segment of the indicated genotypes. *Nmnat^+/-^* mutants exhibit comparable *upd3* levels to controls, but ISR induction in C4da neurons significantly increases *upd3* levels. P values, ANOVA with a post-hoc Šídák’s multiple comparisons test. **(F)** *upd3* is required for LDD. Plot depicts effects of *upd3* mutations (*Δ/+*, heterozygous control; *Δ/0*, hemizygous males; *ΔΔ* homozygous mutant females) on presynaptic gradient measurements in controls, *Nmnat^+/-^* mutants, and larvae with C4da-specific *path^RNAi^* or *PEK* expression. P values, ANOVA with a post-hoc Šídák’s test. **(G-H)** *upd3* is required cell-autonomously in C4da neurons for LDD. **(G)** Plot depicts effects of *upd3* or control RNAi on presynaptic gradient induced by *path* inactivation in C4da neurons. P values, ANOVA with a post-hoc Tukey’s test. **(H)** C4da-specific *upd3* expression suppresses the posterior hyposensitivity to noxious mechanical inputs caused by *path* inactivation. P values, Chi-square test. **(I-J)** *upd3* expression is sufficient to induce LDD phenotypes. **(I)** *upd3* expression in C4da neurons induces a gradient of presynapses that is suppressed by *Tb*, which reduces axon length. P values, ANOVA with a post-hoc Tukey’s test. **(J)** *upd3* expression in C4da neurons induces a gradient of sensitivity to noxious mechanical inputs, with stimulation of anterior segments eliciting a higher frequency of nocifensive rolls compared to posterior segments. P values, Chi-square test. **(K)** Graded effects on presynapses (X-axis, mean of presynaptic gradients) and nocifensive behavior (Y-axis, difference in behavioral response to A and P stimulation) are highly correlated. The line depicts the linear regression to the indicated data points and dashed lines demarcate confidence bands. **(L)** Ectopic *upd3* expression in the VNC induces LDD phenotypes in C4da neurons. Plot depicts measurements of the presynaptic gradient in C4da neurons of control larvae or larvae expressing *UAS-upd3* with the indicated *GAL4* drivers. *upd3* expression in cho neurons or in ABLK interneurons but not in body wall epidermal cells induces LDD. P-values, ANOVA with a post-hoc Dunnett’s test.

We focused our attention on the top differentially expressed gene *unpaired 3 (upd3)*, which encodes one of three *Drosophila* unpaired family cytokines and functions broadly in controlling inflammation and proliferation (*42–47*)(Fig. 2B). Although nervous system functions for *upd3* have not been defined, *upd3* expression is induced by elevated ROS levels in pericardial cells (*48*) and by the zinc-finger transcription factor *Xrp1* in eye imaginal disc cells experiencing ISR (*49*). Furthermore, the *upd3* orthologue IL-6 has pleiotropic functions in the mammalian nervous system including context-dependent contributions to neuronal survival and degeneration (*50*, *51*). Based on our findings that *path* inactivation and ISR induction elevated ROS levels, increased *Xrp1* expression (table S1), and increased *upd3* expression (Fig. 2B), we hypothesized that udp3 functions as a stress-induced cytokine which mediates LDD.

To test this hypothesis, we examined whether *upd3* expression was responsive to conditions that induced LDD, LID, or both. To this end, we used hybridization chain reaction fluorescent mRNA *in situ* hybridization (HCR-FISH) to monitor *upd3* mRNA levels in C4da neurons. First, as a control for probe sensitivity, we monitored *upd3* levels in C4da neurons overexpressing *upd3*, which exhibited robust HCR-FISH signal when stained with *upd3* probes (fig. S6A). Next, we assayed for *upd3* expression in conditions that induced LDD as well as an amorphic *upd3* mutant (*upd3^Δ^*) as a control for specificity. Whereas *upd3* mRNA was present at low levels in controls, *upd3* was significantly upregulated by *path* inactivation or ISR activation in C4da neurons (Fig. 2, C to E, and fig. S6B). We further found that *upd3* was induced to a similar degree in all abdominal segments (Fig. 2E and fig. S6, D to F), suggesting that *upd3* mRNA levels are not directly coupled to axon length. In contrast, the *upd3* mRNA signal that was induced by ISR activation in C4da neurons was absent from homozygous *upd3^Δ^* mutant larvae (fig. S6, F and G). Unlike in conditions inducing LDD, heterozygous loss of *dNmnat*, which triggered LID, did not significantly upregulate *upd3* expression (Fig. 2, D and E, and fig. S6H).

Next, to determine whether *upd3* is necessary for LDD we monitored effects of *upd3* knockout (*upd3^Δ^*) on treatments that induced LDD and/or LID. On its own, *upd3* inactivation had no effect on overall synapse numbers (Fig. 2F and fig. S7A), demonstrating that *upd3* is dispensable for normal development of C4da axons. In contrast, *upd3* mutation suppressed the presynaptic gradient induced by *path* inactivation or ISR activation in C4da neurons (Fig. 2F and fig. S7B). Selective *upd3* inactivation in C4da neurons using RNAi suppressed both the presynaptic gradient and the posterior hyposensitivity to mechanonociceptive stimuli induced by *path* inactivation (Fig. 2, G and H, and fig. S7C), demonstrating that *upd3* is required cell-autonomously in C4da neurons to trigger LDD phenotypes. Finally, we found that *upd3* preferentially affected LDD: *upd3* mutation suppressed LDD phenotypes but not presynapse loss in anterior segments that occurred as a result of *path* inactivation or ISR activation (fig. S7A), and additionally failed to suppress LID phenotypes of *dNmnat^+/-^* mutants (fig. S7A).

To evaluate whether *upd3* expression is sufficient to trigger LDD we overexpressed *upd3* selectively in C4da neurons and assayed for effects on C4da presynapse numbers and nociceptive sensitivity. Indeed, *upd3* overexpression induced a presynaptic gradient that was dependent on axon length without affecting presynapse numbers in anterior segments (Fig. 2I and fig. S7D). As with other conditions that induced LDD, larvae overexpressing *upd3* in C4da neurons exhibited significantly reduced nocifensive responses to posterior compared to anterior stimuli (Fig. 2J); we note that *upd3*-overexpressing larvae exhibited an apparent increase in nocifensive responses to anterior stimuli when compared to driver-only or *UAS-upd3* controls. Overall, we found that the extent of posterior hyposensitivity induced by these treatments was tightly correlated with the severity of the presynaptic gradient (Fig. 2K).

Although *upd3*-dependent LDD phenotypes were suppressed by shortening axons using *Tb*^1^ mutants (Fig. 2I and fig. S7D), we found no evidence that *upd3* mRNA expression was graded in conditions that induced LDD. We therefore hypothesized that factors other than neuronal *upd3* levels account for *upd3* effects on LDD. To test this possibility, we examined whether supplying *upd3* in trans induced LDD. Ectopically expressing *upd3* in chordotonal (cho) neurons, a distinct population of PNS neurons whose axons target a region of the neuropil adjacent to C4da axons, induced a presynaptic gradient in C4da neurons due to a reduction in presynapses in posterior segments (Fig. 2L and fig. S7E).

Hence, even in the absence of signals that trigger ISR in C4da neurons, synapses from C4da neurons with the longest axons are intrinsically vulnerable to synapse loss induced by upd3 signaling. Likewise, ectopic *upd3* expression in abdominal leucokinin (ABLK) neurons, but not in body wall epidermal cells induced LDD phenotypes (Fig. 2L and fig. S7E). ABLK neurons are third-order interneurons in the nociceptive circuit that do not directly synapse with C4da axons (*52*). Taken together, these data are consistent with upd3 locally signaling to cells within the VNC to trigger axon length-dependent presynapse removal from C4da neurons.

### upd3 activates glial cells to remove vulnerable synapses

Like its human orthologue IL-6, upd3 activates the JAK/STAT pathway in target cells by binding to its cognate receptor domeless (dome). Ligand binding to dome induces activation of the Janus tyrosine kinase hopscotch (hop), which phosphorylates dome, creating a docking site for subsequent phosphorylation and activation of Stat92E (Fig. 3A) (*42*, *53*).

**Fig. 3.**
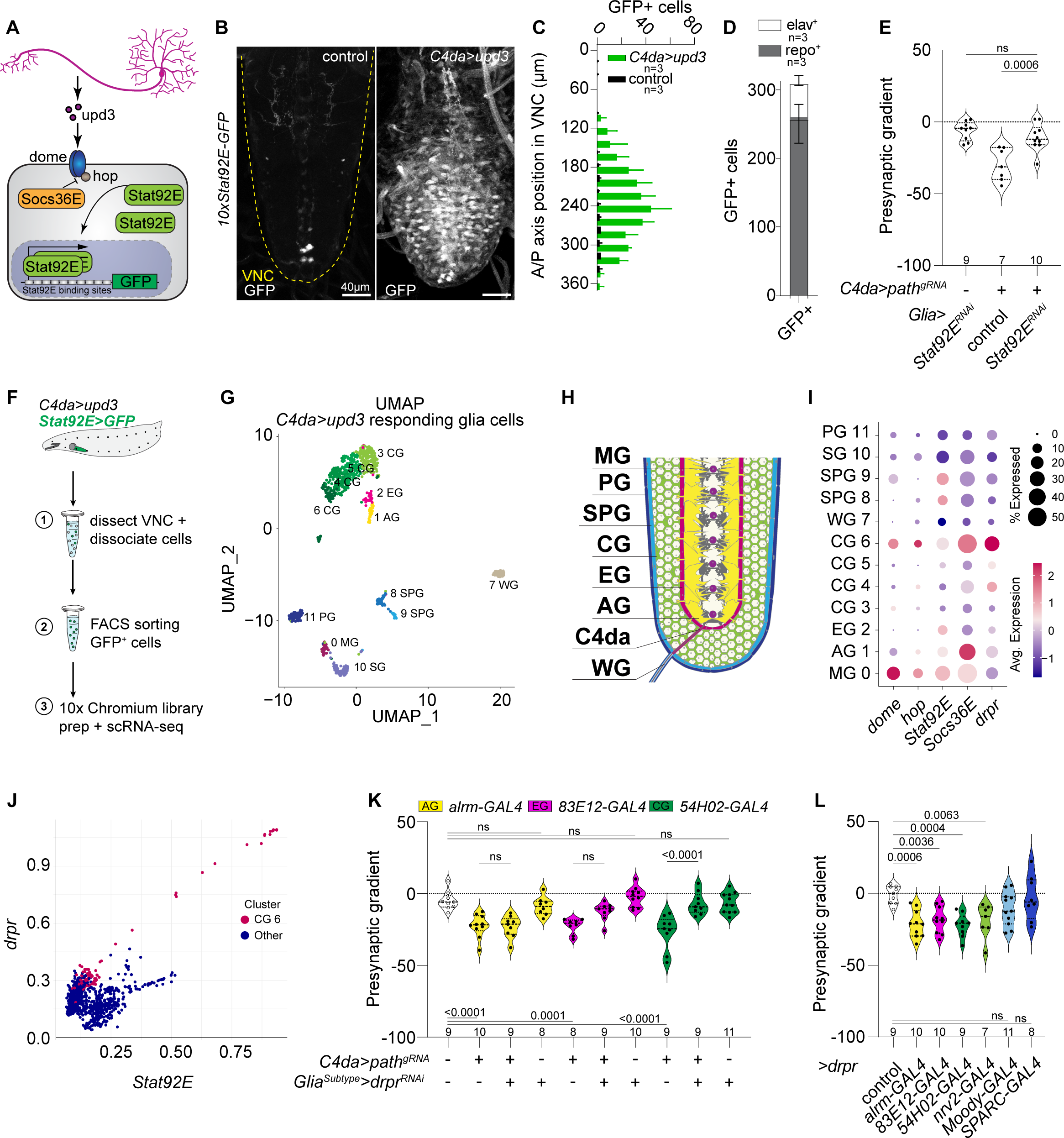
upd3 signals to glia to mediate LDD phenotypes. **(A)** Schematic of upd3 signaling pathway and *10xStat92E-GFP* reporter transgene. **(B-D)** *upd3* expression in C4da neurons triggers widespread Stat92E activity in the VNC. Representative maximum projections of confocal stacks show *10xStat92E-GFP* reporter expression in a control VNC (*ppk-GAL4/+*) and a VNC from a larva expressing upd3 in C4da neurons (*ppk-GAL4, UAS-upd3/+*). **(C)** *Upd3*-responsive cells are concentrated in the posterior portion of the VNC, and antibody staining revealed that these cells are predominantly glia **(D)**. The plot depicts mean and standard deviation values for *10xStat92E-GFP* expressing cells in each VNC (*ppk-GAL4, UAS-upd3/+*) that exhibited elav or repo immunoreactivity. **(E)** Glial *Stat92E* is required for LDD. *Stat92E* knockdown in glia does not induce a presynaptic gradient in C4da neurons or affect C4da neuron presynapse numbers (fig. S9), but it significantly ameliorates LDD triggered by *path* inactivation in C4da neurons. P values, ANOVA with post-hoc Šídák’s test. **(F-G)** Identification of *upd3*-responsive cells in the VNC using scRNA-seq. **(F)** Schematic depicting scRNA-seq sample preparation workflow. **(G)** UMAP plot depicting 12 clusters of *upd3*-responsive glial cells. Glial cells were identified based on marker gene expression (fig. S10B) and include cells from each of the glial types depicted in **(I)**. Abbreviations: AG, astrocytic glia (yellow); CG, cortex glia (green); EG, ensheathing glia (burgundy); MG, midline glia (purple); PG, perineural glia (blue); SPG, subperineural glia (aqua); WG, wrapping glia (light gray). C4da neuron terminal axonal arbors are depicted in gray. **(J-K)** Expression of JAK/STAT pathway genes in *upd3*-responsive cell clusters. **(J)** JAK/STAT pathway genes are most highly expressed in a cluster of cortex glia (cluster 6) and midline glia (cluster 0), with cluster 6 exhibiting high levels of *drpr* expression. **(K)** Among all *upd3*-responsive glial cells, a subset of cluster 6 cells exhibits the highest levels of *Stat92E* and *drpr* expression. **(L)** *drpr* is required in cortex glia for LDD. RNAi of *drpr* in CG but not AG, EG, SPG, or PG significantly ameliorates presynaptic gradient induced by *path* inactivation in C4da neurons (see also fig. S11A). P-values, Kruskal-Wallis test with post-hoc Dunn’s test. **(M)** Overexpressing *drpr* in AG, CG, or EG is sufficient to induce LDD. P values, ANOVA with post-hoc Dunnett’s test.

Stat92E acts as transcriptional activator for the inhibitory factor *Socs36E*, therefore we used a GFP reporter containing multimerized Stat92E binding sites derived from the *Socs32E* enhancer (*10xStat92E-GFP*) as a proxy for Stat92E transcriptional activity (Fig. 3A)(*54*, *55*). First, we assayed for Stat92E activity in wild-type control third instar larvae, which exhibited minimal reporter activity in the VNC (Fig. 3B). In contrast, *upd3* overexpression in C4da neurons induced broad *10xStat92E-GFP* expression within the VNC, with the majority of these GFP^+^ cells located in in the posterior ventral portion of the VNC adjacent to C4da axon termini (Fig. 3, B and C, and fig. S8, A to D).

Antibody staining revealed that most of the *10xStat92E-GFP*-expressing cells also expressed the pan-glial marker repo (Fig. 3, D and E; fig. S8, A to D), therefore we next examined whether upd3 activation of glial cells contributes to LDD. To test this possibility, we assayed effects of glial JAK/STAT pathway inactivation on *path* mutant LDD phenotypes in C4da neurons. Indeed, glia-specific RNAi knockdown of *Stat92E* significantly attenuated the synaptic gradient induced by *path* inactivation (Fig. 3E and fig. S9A). Furthermore, overexpressing the JAK/STAT pathway activator *hop* in subsets of glial cells including subperineurial glia (SPG), astrocyte-like glia (AG) and cortex glia (CG) induced LDD but not LID (fig. S9, B and C). Taken together, our results demonstrate that JAK/STAT pathway activation in glia cells is necessary and sufficient for LDD.

### Stat92E activity correlates with engulfment receptor draper in subsets of glia cells

Gut-derived upd3 signaling mediates metabolic reprogramming of ensheathing glia (EG) upon enteropathogen infection (*56*). We therefore hypothesized that SSN-derived upd3 activates glial cells to trigger presynapse engulfment. Glial phagocytic clearance of injured axons mediated by the MEGF10 family engulfment receptor draper (drpr) is dependent on Stat92E activity (*56–60*), therefore as a first test of our model we assayed requirements for glial *drpr* in LDD. We found that selective knockout of *drpr* in glial cells using the CRISPR/Cas9 system abolished the presynaptic gradient induced by C4da neuron-specific inactivation of *path*, inactivation of *ppp1R15,* or overexpression of *upd3* without affecting the number of presynapses in anterior segments (fig. S9, D and E), demonstrating that *drpr* is required in glial cells for LDD triggered by upd3.

To systematically identify the glial cells responsive to upd3 cytokine from C4da neurons we overexpressed *upd3* in C4da neurons, dissociated larval VNCs expressing the *10xStat92E-GFP* reporter, and subjected FACS-sorted GFP+ cells to single-cell RNA sequencing using droplet microfluidics (Fig. 3F). After stringent quality-based filtering we used graph-based methods to cluster 1701 cells based on shared gene expression programs (*61*). This initial clustering analysis yielded 18 cell clusters (fig. S10A) which were evaluated according to marker gene expression, revealing 10 clusters containing 913 cells that expressed glial markers. We isolated and reclustered these glial cells, revealing 12 glial clusters that exhibited *10xStat92E-GFP* reporter activity in response to C4da neuron *upd3* expression, prominently included CG (4 clusters, 58.4% of cells) as well as SPG (2 clusters), AG, EG, midline glia (MG), perineural glia (PG), surface glia (SG), and wrapping glia (WG) (Fig. 3, G and H, and fig. S10B).

To identify the glial subsets most likely to mediate C4da presynapse removal, we queried for expression of JAK/STAT pathway components including *dome*, *hop*, *Stat92E,* and the Stat92E transcriptional targets *Socs36E* and *drpr*. We found that two clusters, corresponding to CG and MG, exhibited high expression of JAK/STAT pathway components; among these, only the CG exhibited high levels of *drpr* expression (Fig. 3I). Furthermore, we found that expression of *drpr* and *Stat92E* was highly correlated in a subset of CG but not other glial subtypes (Fig. 3J and fig. S10C), therefore we hypothesized that these CG cells were responsible for *upd3*-dependent engulfment of C4da presynapses.

### Cortex glia utilize draper to target presynapses in LDD

To assay contributions of glial subtypes to LDD, we induced LDD by inactivating *path* in C4da neurons and additionally knocked down *drpr* (*drpr^RNAi^*) using *GAL4* driver lines that target distinct glial subtypes. The PG and SPG cells in our scRNAseq dataset exhibited low levels of *drpr* expression, and we found that *drpr* knockdown using drivers for PG (*SPARC-GAL4*) or SPG (*Moody-GAL4*) had no significant effect on LDD induced by *path* inactivation (fig. S11, A and B). Prior studies defined functions for AG, CG and EG in phagocytic engulfment, but we found that *drpr* knockdown using the EG driver *83E12-GAL4* or the AG driver *alrm-GAL4* had no significant effects on LDD (Fig. 3K and fig. S11B). In contrast, LDD was significantly suppressed by expressing *drpr* RNAi with the CG driver *54H02-GAL4*, and we corroborated this result with an additional CG driver (*nrv-2-GAL4*) (Fig. 3K and fig. S11, A and B). Prior studies defined a role for insulin receptor (InR) in *drpr*-mediated phagocytic activity following axotomy (*57*), however glial *InR* inactivation had no effect on LDD in C4da neurons (fig. S11, C and D), suggesting that injury-induced phagocytic clearance by EG and *upd3*-dependent phagocytic activity in LDD involve distinct regulatory mechanisms.

Next, we examined the capacity of different glial cells to induce LDD. To this end, we overexpressed *drpr* in glial cells and monitored C4da neuron presynapse numbers. We found that overexpressing *drpr* in AG, CG, or EG, but not other glial types induced LDD in an otherwise wild-type background (Fig. 3L and fig. S11E). These data demonstrate that presynapses of C4da neurons innervating posterior segments were either selectively vulnerable for removal by these glial subtypes or that these glia subtypes were selectively primed to remove synapses in posterior segments. Among these treatments, *drpr* overexpression in AG but no other glial subtypes additionally reduced presynapse numbers in anterior segments (fig. S11E), hence although AG exhibit heightened activity towards C4da presynapses on long axons, they exhibited less selectivity than other glial subtypes.

### Fly models of NDDs induce upd3-dependent LDD

Selective vulnerability of neurons with extensive projections and graded axon-length dependent degeneration are pathological features of many NDDs. We, therefore, examined whether fly models of human NDDs induced LDD in C4da neurons and, if so, if the length-dependent phenotypes involved *upd3*. We assayed for LDD phenotypes in larvae in which we either targeted *Drosophila* orthologues of genes associated with NDDs including human *SOD2 (AD/PD/ALS*(*62–65*)) and *PRKN* (PD(*66–69*)), or overexpressed transgenes containing dominant disease-associated mutations including *α-Synuclein hSNCA^E46K^* (PD(*70–73*)) and *ATXN3* with a glutamine tract expansion (*hATXN3^.tr-Q78^*; SCA3/polyQ (*74–77*)). First, we assessed functional consequences of the various treatments and found that each treatment significantly reduced nocifensive responses to mechanosensory stimuli delivered to posterior but not anterior segments, a hallmark of LDD in C4da neurons (Fig. 4, A and B).

**Fig. 4.**
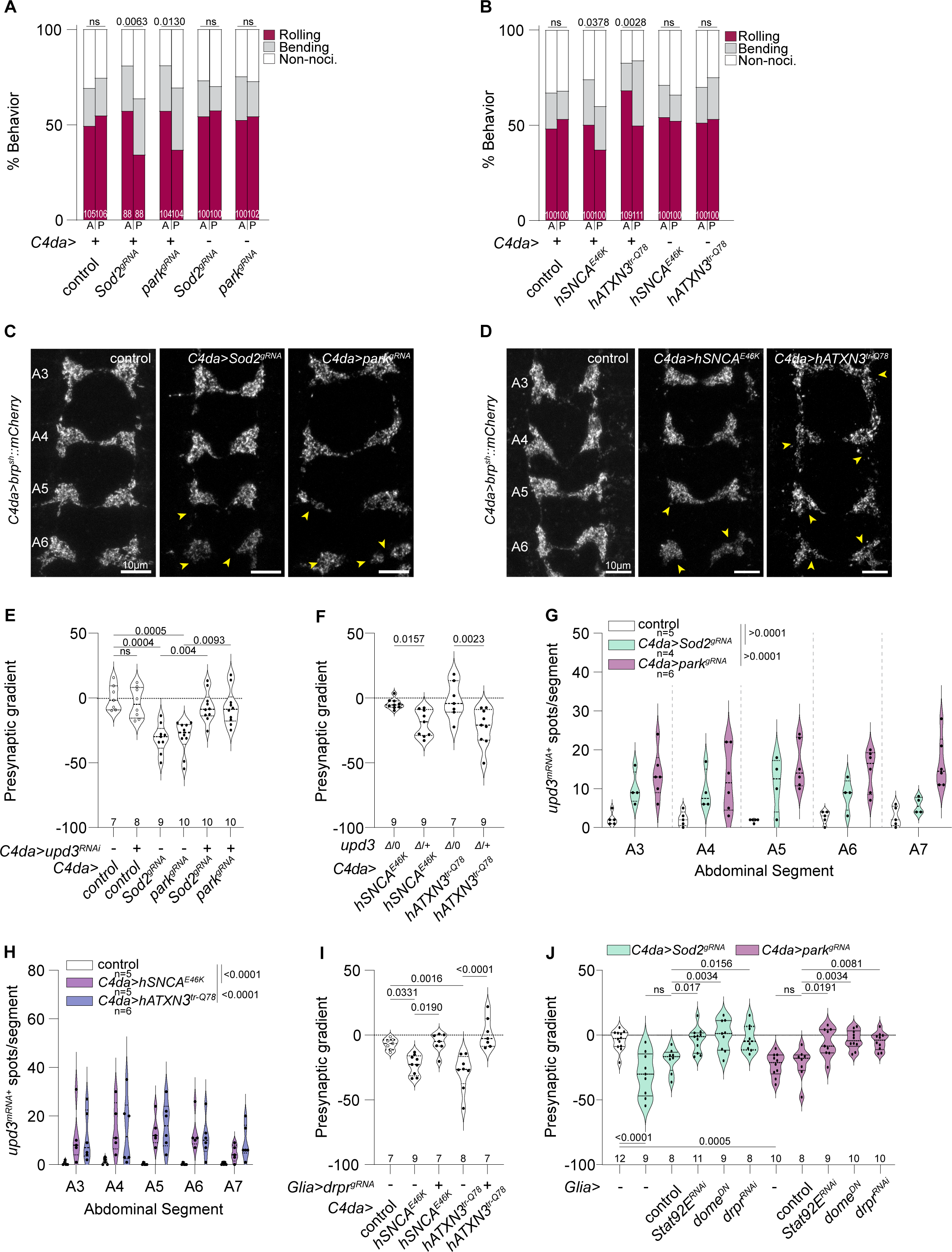
*Drosophila* models of NDDs exhibit *upd-3* dependent LDD. **(A)** Inactivation of *Sod2* or *park* in C4da neurons induces a gradient of sensitivity to noxious mechanical inputs. Plot depicts the proportion of larvae of the indicated genotypes which exhibit rolling (red), bending (gray), or non-nocifensive responses to noxious mechanical stimuli (see Fig. 1J). P values, Chi-square test compared to effector only control. **(B)** Overexpression of *hATXN3^tr.Q78^ or hSNCA^E46K^* in C4da neurons induce gradients of sensitivity to noxious mechanical inputs. Plot depicts the proportion of larvae of the indicated genotypes which exhibit rolling (red), bending (gray), or non-nocifensive responses to noxious mechanical stimuli (see Fig. 1J). P values, Chi-square test compared to effector only control. **(C)** Maximum intensity projections depict the distribution of C4da neuron presynapses labeled by *ppk::brp^sh^-mCherry* in the VNC Segments A3-A6 in larvae with *Sod2* or *park* inactivation in C4da neurons. Arrows (yellow) indicate sites of strong synaptic loss. **(D)** Maximum intensity projections depict the distribution of C4da neuron presynapses labeled by *ppk::brp^sh^-mCherry* in the VNC Segments A3-A6 in larvae with *hATXN3^tr.Q78^ or hSNCA^E46K^* overexpression in C4da neurons. Arrows (yellow) indicate sites of strong synaptic loss. **(E)** Violin plot depicts presynaptic gradient of *Sod2* or *park* inactivation in C4da neurons with and without simultaneous RNAi against *upd3*. *Sod2* or *park* inactivation drives robust LDD, which is abolished by *upd3* knockdown. P-Values, ANOVA with post-hoc Šídák’s multiple comparison test. **(F)** Violin plot depicts presynaptic gradient of *hATXN3^tr.Q78^ or hSNCA^E46K^* overexpression in C4da neurons in hetero- or hemizygous mutant background for *upd3*. Significant differences in presynaptic gradient in *upd3Δ* hemizygous compared to homozygous background during overexpression of *hATXN3^tr-Q78^ or hSNCA^E46K^* in C4da neurons. P-Values, ANOVA with post-hoc Šídák’s multiple comparison test. **(G-H)** *upd3* mRNA expression is induced in C4da neurons undergoing NDD models. **(G)** Violin plots depict the number of *upd3* puncta with intensity > 8000 AU in each abdominal segment of the indicated genotypes. *Sod2* or *park* inactivation exhibit significantly increased *upd3* levels. P values, Two-Way ANOVA with a post-hoc Dunnett’s multiple comparisons test. **(H)** Violin plots depict the number of *upd3* puncta with intensity > 8000 AU in each abdominal segment of the indicated genotypes. *hATXN3^tr.Q78^ or hSNCA^E46K^* overexpression induces significantly increased *upd3* levels. P values, Two-Way ANOVA with a post-hoc Dunnett’s multiple comparisons test. **(I)** Inactivation of *drpr* in glia cells significantly ameliorates presynaptic gradient induced by *Sod2* or *park* inactivation in C4da neurons. P-Values, ANOVA with post-hoc Šídák’s multiple comparison test. **(J)** Inactivation of *Stat92E,* overexpression of DN version of upd3 receptor *dome,* or knockdown of *drpr* in glia cells abolishes presynaptic gradient induced by *Sod2* or *park* inactivation in C4da neurons. P-Values, ANOVA with post-hoc Šídák’s multiple comparison test.

Furthermore, although the various treatments induced degenerative phenotypes with distinctive features (Fig. 4, C and D), each treatment preferentially affected presynapses of C4da neurons innervating posterior segments (Fig. 4, E and F; fig. S12, A and B). All NDD models induced *upd3* expression in C4da neurons (Fig. 4, G and H; fig. S13, A and B), and in each case the gradient of presynapse loss was significantly ameliorated by *upd3* inactivation (Fig. 4, E and F), suggesting that *upd3* is a nexus for LDD signaling. Finally, we confirmed that LDD triggered by each perturbation was dependent on *drpr* function and JAK/STAT signaling in glia cells (Fig. 4, I and J, and fig. S14, A and B). Altogether, these studies demonstrate that axon length-dependent presynapse loss triggered by a variety of NDD-related genetic programs involves *upd3*-dependent recruitment of phagocytic glia.

## DISCUSSION

Selective vulnerability of neurons with extensive projections and graded axon-length dependent degeneration are shared pathological features of many NDDs, yet the basis for these length-dependent effects is poorly understood. Here, we describe a new experimental system for quantitative genetic dissection and behavioral analysis of axon length-dependent degeneration. Axons of C4da neurons that innervate posterior segments grow more than those innervating anterior segments while forming equivalent numbers of synaptic contacts, and we find evidence that under degenerative conditions this model recapitulates many hallmarks of selective vulnerability in NDDs (*78*). This includes defects in proteostasis inducing the ISR, redox imbalance, synaptic network dysfunction and inflammatory responses. The evolutionary conserved ISR is linked to numerous NDDs including PD, ALS, AD and CMT (*37*, *38*) and, for example, has been shown to contribute to selective vulnerability of subsets of upper motor neurons in ALS mouse models (*79*). Proteostasis defects due to protein misfolding and aggregation or oxidative stress activate different stress sensor kinases including Pkr, Gcn2 or Perk, resulting in eIF2α phosphorylation and consequential shutdown of general translation (*80*). In turn, the ISR selectively upregulates Atf4 and CCAAT Enhancer Binding Protein Gamma (C/EBP-γ)-dependent transcriptional programs, which can trigger the cell’s innate immune response including upregulation of the inflammatory cytokine IL-6 (*37*, *81*, *82*) . Similarly, Xrp1, a homolog of mammalian C/EBPs (*83*), triggers ISR mediated stress response programs, including upd3 induction(*49*). In C4da neurons, ISR activation drives progressive loss of presynapses and deficits in mechanonociception that covaried in severity with increasing axon length. Moreover, ISR activation induced neuronal expression of the IL-6 orthologue upd3, which we identified as an evolutionarily conserved central mediator of LDD. Expression of *upd3* selectively affects LDD but not LID and was not induced by Wallerian degeneration-linked perturbations, demonstrating that the two phenomena are genetically separable. Remarkably, *upd3* expression in other neuronal subsets could trigger non-cell autonomous LDD in C4da neurons. Since the degeneration of specific neuronal subsets which eventually spreads to other populations is another hallmark of NDDs (*2*), our results suggest that inflammatory cytokine signaling could in part mediate this spreading. Axonal spreading of pathology is discussed as a potential disease spread mechanism in PD and ALS (*84*, *85*) and could be triggered by guiding microglia to large axon projections from other brain regions. Notably, inflammation and inflammatory microglia are suggested to exacerbate neurodegenerative progression and prion-like seeding (*86–88*).

Chronic inflammation is observed in most NDDs (*3*, *89*, *90*) and IL-6 levels were reported to be elevated in AD, PD and ALS patients (*50*, *51*). IL-6 levels positively correlate with cognitive impairment in AD patients and mouse models (*91*), and increased levels of IL-6 have been linked to poor disease prognosis in PD (*92*). Inflammation is also a key feature in PD (*93*), yet the sequence of events of how inflammatory processes are first induced and precisely how they contribute to neurodegenerative processes is not well understood. ISR activation and general inflammatory processes in the brain can upregulate IL-6 expression in neurons as well as astrocytes and microglia, causing a proinflammatory cascade that can drive neurodegeneration through the IL-6 mediated activation of the JAK/STAT pathway (*94–96*). IL-6 and chronic inflammation can further lead to increased permeability of the blood brain barrier and invasion of immune cells into the CNS, further promoting neurodegeneration (*94*, *97*). Our in vivo studies suggest that upd3/IL-6 drives glial activation and phagocytosis of axon terminals. In line with this notion, neuronal IL-6 is elevated upon neuronal injury (*98*, *99*), and injured peripheral neurons induce STAT3-dependent microglia activation in the mouse spinal cord (*100*).

Our studies reveal that presynapses on neurons with long axons are more vulnerable to removal than those on same-type neurons with short axons. What is the cause of axon length-dependent synapse loss? We found that LDD phenotypes were dependent on the inflammatory cytokine upd3, yet we found no evidence that *upd3* mRNA expression was graded in conditions that induced LDD. Although we cannot rule out contributions of post-transcriptional control of upd3 levels, three key pieces of evidence suggest that additional factors contribute to presynapse vulnerability: providing *upd3* in trans generated LDD phenotypes, axon shortening suppressed LDD induced by *upd3* expression, and *drpr* overexpression in glial cells induced LDD. While defects in axonal transport have been extensively linked to early events of degeneration in NDDs (*101*, *102*), LDD can be triggered by ectopic *upd3* expression or activation of glial JAK/STAT signaling in an otherwise wild-type background. This suggests that presynapses of unaffected neurons with long axons are intrinsically more vulnerable to synapse loss, which was also corroborated by reduced axon length suppressing LDD.

One possible source of this vulnerability is the graded expression of a neuronal “eat me” signal that promotes preferential presynapse removal from neurons with long axons. Our finding that presynapse loss in LDD is drpr-dependent and mediated by a subset of glial cells suggests that drpr and/or other glial cell surface molecules mediate molecular recognition of presynapses marked for removal. Of note, the most extensively characterized signal that promotes phagocytic engulfment, phosphatidyl serine (PS) exposure, drives *drpr*-dependent phagocytosis of degenerating neurites in *Drosophila* (*103*, *104*). Under normal conditions PS remains largely confined to the inner leaflet of the plasma membrane, and this PS asymmetry is achieved through the activity of Type IV P-type-ATPases, also known as flippases (*105*), which additionally require a CDC50A/TMEM30A family chaperone for their correct localization and function. Hence, alterations in flippase activity during degenerative conditions could expose PS and contribute to LDD. Indeed, inactivation of flippase complex component *ATP8A1* drives PS externalization in hippocampal neurons, mutations of ATPA2 are linked to a rare form of cerebellar ataxia (*106*, *107*), and Cdc50a knockdown leads to phosphatidylserine exposure at synapses (*108*). Furthermore, Aβ oligomer stimulation of neuronal spines drive PS externalization, triggering TREM2 mediated engulfment by microglia in an AD model (*109*), a mechanism similar to PS-mediated developmental synaptic pruning (*110*). Alternatively, the increased demands for transport in long axons may limit trafficking of flippases and accessory factors to presynapses, limiting the ability of neurons to maintain PS asymmetry.

In mammals, PS exposure can recruit the complement protein C1q, which prompts synapse pruning mediated by a variety of receptors (*111*, *112*). Intriguingly, activation of the innate immune system, including the mammalian complement system (CS), has been implicated in excessive synaptic pruning in AD (*113–117*). CS components are found at pre- and postsynaptic sites where they may act as potential synaptic pruning tags for microglia driven phagocytosis. Complement-related genes are conserved in *Drosophila* and the complement-related gene *mcr* is required for *drpr*-dependent autophagy of neighboring cells at sites of injury (*118*). Hence, future studies in Drosophila could reveal connections between axon length, maintenance of PS asymmetry, and LDD.

In addition to PS, *Drosophila* drpr recognizes protein ligands that serve as damage-associated molecular patterns (DAMPs) and facilitate apoptotic engulfment in certain contexts (*119*, *120*). Glial phagocytic receptors for DAMPs have been implicated in a variety of NDDs, notably including the AD risk gene TREM2 which can modulate complement-mediated synapse loss through binding to C1q (*116*). Hence graded expression of DAMPs or gradations in receptor activity could likewise contribute to LDD.

Our studies provide evidence that neuronal complexity, i.e. axon length, is a deterministic factor in the vulnerability of C4da axons to degeneration. First, presynapse loss phenotypes appear earliest and are most severe in C4da neurons that innervate posterior segments and hence have the longest axons. Second, treatments that induce LDD but not LID trigger presynapse loss selectively in neurons with long axons. Third, shortening axons suppressed whereas lengthening axons enhanced LDD phenotypes. As both loss and gain of function of several genes linked to PD, ALS, and SCA3/polyQ disease resulted in LDD of C4da axons this suggests a conserved and converging mechanism shared by multiple NDDs, opening the possibility that neuron-driven cytokine-mediated phagocytosis is a common feature of different neurodegenerative pathways resulting in LDD.

## Acknowledgements

Fly stocks obtained from the Bloomington Drosophila Stock Center (NIHP40OD018537) and antibodies obtained from the Developmental Studies Hybridoma bank, created by the NICHD of the NIH and maintained at The University of Iowa, were used in this study. We thank Meike Petersen for cloning of ppk-Brp^sh^-mCherry, David Bilder and Hyung Don Roo for sharing fly strains, and the Rasmussen lab for sharing expertise and reagents for HCR-FISH.

## Author Contributions

Conceptualization: FT, PS, and JZP

Methodology: FT and CY

Investigation: FT, CY, JH, and ND

Data curation: FT and CY

Formal Analysis: FT and CY

Visualization: FT Supervision: PS and JZP

Funding acquisition: FT, PS, and JZP

Writing – original draft: FT, PS, and JZP

Writing – review and editing: FT, PS, and JZP

## Supplementary Materials

Materials and Methods

Figs. S1 to S14

Tables S1 to S4

Supplementary References

