## Supplemental Figures and Methods for "Axon length-dependent synapse loss is mediated by neuronal cytokine-induced glial phagocytosis"

### Supplementary Materials

#### Materials and Methods

##### *Drosophila* cultures

All fly stocks were kept at 25°C and 70% relative humidity with a 12 h light/dark cycle on standard cornmeal-molasses fly food. Stocks were obtained from the Bloomington Drosophila Stock Center (BDSC) unless otherwise indicated. A complete list of alleles used in this study is provided in table S2.

##### *ppk-brp$^{sh}$::mCherry* fusion construct

brp$^{sh}$-mCherry was cloned into the pENTR vector and transferred into a pDEST vector containing the ppk enhancer (PMID: 21606367) using the GATEWAY system (Thermo Fisher Scientific, Carlsbad, CA). Transgenic flies were generated by inserting the attB containing pDEST-brp$^{sh}$-mCherry plasmid into the 3$^{rd}$ chromosome of flies containing an attP landing site (VK27, BDSC #9744) using phiC31 mediated transgenesis (PMID: 15126397).

##### Confocal microscopy and immunohistochemistry

Specimens (larval fillet preparations or isolated larval brains) were dissected in PBS and fixed in PBS containing 4% paraformaldehyde for 15 min at room temperature. Samples were rinsed 3 x 1 min in PBS and processed for direct imaging of native fluorescence or immunohistochemistry. To image native fluorescence, brains were mounted on coverslips coated with poly-L-lysine (Millipore-Sigma, St. Louis, MO) in SlowFade Gold mounting medium (Thermo Fisher Scientific, Carlsbad, CA) and imaged using confocal microscopy (Leica SP5, Zeiss LSM 900). Confocal Z-Stacks were analyzed and processed in Fiji (ImageJ, NIH, Bethesda) and/or Imaris (BitPlane, Belfast, UK). For visualizing structures with immunohistochemistry, fixed specimens were washed three times with PBS + 0.3% Triton X-100 (PBSTx), incubated in blocking buffer (5% NGS in PBS) for 30 min with subsequent incubation of primary antibody (4° C overnight). After 3 x 5 min washing with PBSTx , samples were stained with fluorescently conjugated secondary antibodies diluted in 5% NGS for 1 h at room temperature and washed 3 x 5 min with PBSTx then once with PBS prior to mounting. Refer to table S2 for a detailed list of antibodies and dilutions.

##### Presynapse quantification

Presynaptic active zones of C4da neurons were labeled by brp$^{short}$::mCherry and imaged in confocal Z-stacks with a step size of 250 nm. Confocal stacks were processed and analyzed in Imaris (Bitplane AG, Zurich, Switzerland) as follows. First, images were batch-processed with baseline subtraction and Clear View deconvolution. Next, presynaptic puncta of C4da neurons in A3-A6 segmental domains were semi-automatically detected using the spot function (200 nm with Z-Axis correction). Fixed intensity quality thresholds were used, based on automatic detection thresholds for each set. Presynaptic gradients were defined by the slope (b) of a linear regression curve through presynaptic puncta (dependent variable y) from abdominal segments 3 to 6 (independent variable x) for each animal.

**Mechanonociception assay**

Reaction to noxious mechanical force was assayed with 50 mN calibrated *von-Frey*-filaments using tightly staged third instar larvae (96 ± 3 h AEL). Larvae were isolated from media, washed in distilled water, and transferred with a paintbrush to a moistened 2% agar plate. One set of animals was stimulated twice within 2 sec on abdominal segments 2-3 (anterior stimulation) and another set was stimulated twice within 2 sec on abdominal segments 6-7 (posterior stimulation). Behavioral responses (non-nociceptive, bending, rolling/multiple rolling) to both stimuli were noted but only behavioral responses to the second stimulus were analyzed and plotted. Staging and behavior assays were done in a blinded and randomized fashion.

**ISRIB feeding**

ISRIB (MilliporeSigma, St. Louis, MO) was dissolved in DMSO and stored at -20°C. ISRIB stock solutions were diluted in water and mixed with inactivated dry yeast to create a yeast paste. Yeast paste with ISRIB or DMSO (vehicle control) was applied to simple grape agar plates. Experimental crosses were staged on normal agar plates and 24 h AEL larvae were transferred onto ISRIB or DMSO plates.

**HCR RNA-FISH**

To quantify *upd3* RNA levels using HCR RNA-FISH (*1*), we adapted a protocol developed for *Drosophila* whole mount brain preparations (dx.doi.org/10.17504/protocols.io.bzh5p386) using DNA probes targeting *upd3* RNA which were designed with the python package "insitu-probe-generator" (*2*). Larval fillet preparations were fixed for 15 min in 4% paraformaldehyde/PBS, washed 3 x 10 min in PBS + 0.3% Triton X-100 (PBSTx), and then incubated with pooled probes (40 probes, 50 pmol each; table S2) (Integrated DNA Technologies, Coralville, IA) for 24 h at 37 °C at 700 rpm in a Thermomixer. Probes were labeled with Alexa Fluor 647 conjugated B3 hairpins (Molecular Instruments, Los Angeles, CA). Samples were additionally immunolabeled with chicken-α-GFP primary antibodies and α-chicken-Alexa Fluor 488 secondary antibodies together with Hoechst 33342 as described above. Confocal Z-Stacks of soma were acquired and spots within somas were counted using Imaris spots function with further analysis and visualization in R/RStudio and Graphpad Prism (Graphpad, San Diego, CA).

**RNA isolation for bulk RNA-seq analysis**

3rd instar larvae with cytoplasmic GFP expressed in C4da were dissected and dissociated in collagenase type I (Thermo Fisher, Bothel, WA) into single cell suspensions as previously described (*3*), with the addition of 1% BSA (MilliporeSigma, St. Louis, WA) to the dissociation mix. After dissociation, cells were transferred to a new 35 mm petri dish with 1 mL 50% Schneider's media (Thermo Fisher, Bothel, WA), 50% PBS supplemented with 1% BSA. Under a fluorescent stereoscope, individual fluorescent cells were manually aspirated with a glass pipette into PBS containing 0.5% BSA, and serially transferred until isolated from cellular debris. Ten cells per sample were pooled, transferred to a mini-well containing 3 µL lysis solution (0.2% Triton X-100 in milliQ water containing 2 U / µL RNAse Inhibitor), lysed by pipetting up and down several times, transferred to a microtube, and stored at -80° C. For the picked cells, 2.3 µL of lysis solution was used as input for library preparation.

**Bulk RNA-seq library preparation**

RNA-Seq libraries were prepared from the picked cells following the Smart-Seq2 protocol for full-length transcriptomes (*4*). To minimize batch effects, primers, enzymes, and buffers were all used from the same lots for all libraries. Libraries were multiplexed, pooled, and purified using AMPure XP beads (Beckman Coulter, Brea, CA), quality was checked on an Agilent TapeStation (Agilent Technologies, Santa Clara, CA), and libraries were sequenced as 51 bp single end reads on a HiSeq4000 at the UCSF Center for Advanced Technology.

**Bulk RNA-seq data analysis**

Reads were demultiplexed with CASAVA (Illumina, San Diego, CA) and read quality was assessed using FastQC (*5*) and MultiQC(*6*). Reads containing adapters were removed using Cutadapt version 2.4 (*7*) and reads were mapped to the *D. melanogaster* transcriptome, FlyBase genome release 6.29, using Kallisto (version 0.46.0) (*8*) with default parameters. Samples were removed from further analysis based on the PCA analysis. Downstream DE analysis was performed by DESeq2 in R. Raw sequencing reads and gene expression estimates are available in the NCBI Sequence Read Archive (SRA) and in the Gene Expression Omnibus (GEO) under accession numbers PRJNA1106555 and PRJNA1106562.

**Sample preparation for single cell RNA-seq**

Staged 3rd instar larvae were sorted under a fluorescent stereomicroscope to confirm *10xStat92E-GFP* reporter expression. VNCs without brain lobes were isolated from the larvae and transferred to 1.5 ml Eppendorf tubes containing dissociation buffer (450 µl PBS supplemented with 1% BSA solution (MilliporeSigma, St. Louis, WA) and 1 mg/mL collagenase type I (Gibco, Thermo Fisher, Bothel, WA). 20 VNCs were dissociated in each of the two sample tubes at 37 °C with mechanical agitation (mixing at 1000 rpm with manual trituration every 10 min). Dissociated cell suspensions were filtered through 70 µm nylon filters to remove cell debris and subsequently stained with propidium iodide (PI) to label dead cells. GFP positive/PI negative neurons were captured using a FACSArial II (BD Biosciences, San Jose, CA). 51,085 and 30,679 events from sample 1 and sample 2 separately have been collected and resuspended in 50 µl PBS + 1% BSA solution after centrifuging for 10 min at 300 x g and 4 °C.

**Library construction and sequencing**

Library construction was performed using a Chromium Single Cell 3' LT v3.1 kit (10X Genomics, Pleasanton, CA). The single-cell suspension was diluted to ~600 cells/µl and loaded onto the chip to target a recovery of approximately 1,000 cells. Following RT-PCR, GEMs were stored at 4 °C overnight followed by cDNA amplification, fragmentation, and indexing the next day. Library quality was evaluated on an Agilent TapeStation (size range of reads from 300 bp to 800 bp with an average size of 500 bp). Sequencing was performed on an Illumina NovaSeq X 10B 100 Cycle Flow Cell by the UCSF, CatCore.

**Computational analysis for scRNA-seq**

*Mapping and counting*

A customized *Drosophila* reference for mapping was generated using the CellRanger (version 6.1.1) 'mkref' function, and 95.7% of reads could be mapped to the Drosophila melanogaster genome (dmel_r6.46_FB2022_03 on FlyBase) after sequencing. A 10X Genomics expression matrix with 666,273,577 reads in total and 319,709 average reads per cell from sample 1 and 509,344,599 reads in total and 336,645 average reads per cell from sample 2 were obtained after mapping and counting by CellRanger (version 6.1.1).

*Quality Control*

The following analyses were performed on R and RStudio, following quality control parameters used in a comparable dataset from Drosophila larvae (*9, 10*). For quality control cells that had fewer than 500 genes, 1000 UMIs, and log (genes) / log (UMIs) ratios of less than 0.8 were removed. Cells with a mitochondrial gene proportion greater than 18%, ribosomal gene proportion less than 5% or greater than 40%, and heat shock gene proportion greater than 5% were removed. We also removed genes with counts less than 2 and expressed in less than 2 cells. Cell multiplets were identified by DoubletFinder (version 2.0.3) and removed.

*Data analysis with Seurat*

1,057 single cells with 11,123 genes expressed for sample 1 and 819 single cells with 11,188 genes expressed for sample 2 were kept as the original dataset. Seurat package (version 4.3.0) in R was used for the following analyses. 2,000 highly variable features were identified through the 'FindVariableFeatures' function and the dataset was scaled by the 'ScaleData' function. To determine the dimensionality of the dataset, we first performed Principal Component Analysis (PCA) and then used the Elbow-Plot and JackStraw-Plot tests together with an evaluation of PC-heatmaps for selecting proper principal components (PCs). 30 PCs were considered to explain the variance of the datasets. Non-linear dimensional reduction was performed by using the 'RunUMAP' function and followed by finding clusters through the 'FindClusters' function with the resolution set at 3.0 for sample 1 and 2.0 for sample 2. The clustering result was visualized in the UMAP plot with 19 clusters (cluster 0 to cluster 18) for sample 1 and 14 clusters (cluster 0 to cluster 13) for sample 2. The average number of total UMIs, number of genes, and sample sizes were calculated at the cluster level. Clusters with lower numbers of total UMIs, number of genes, and sample sizes were considered as heterogenous clusters. After removing heterogenous cells, the filtered dataset contained 924 single cells for sample 1 and 777 single cells for sample 2. Two single-cell datasets were merged into one containing 1701 single cells with 11911 genes expressed using the Seurat 'IntegrateData' function.

The same analysis pipeline was used for the integrated dataset. 22 PCs were considered to explain the variance of the dataset. Non-linear dimensional reduction was performed by using the 'RunUMAP' function and followed by finding clusters through the 'FindClusters' function with the resolution set at 1.0. The clustering result was visualized in the UMAP plot with 18 clusters. Markers of each cluster were defined using the 'FindAllMarkers' function by selecting genes positively expressed in the cluster with the minimum percentage set to 0.5 and log fold change threshold as 0.32. In this way, we identified 2,045 genes as cluster markers for this atlas and used them to annotate the identities of clusters. For gene correlation analysis we used the MAGIC (Markov Affinity-based Graph Imputation of Cells) package by Krishnaswamy lab for imputation before correlation plotting.

**Quantification and statistical analysis**

Statistical analysis was performed with Prism 10 (Graphpad, San Diego, CA) and R/RStudio. All datasets were tested for normality. For normally distributed data, One-Way or Two-Way ANOVA was used, depending on the number of independent variables. For sets including non-normal distributed data, non-parametric Kruskal-Wallis tests were performed. A vs P behavioral data was analyzed using Chi-square tests. Post-hoc tests to correct for multiple comparisons were chosen according to ANOVA design and are indicated in figure legends. P values are indicated in figures, whiskers in plots represent standard deviation, and details of statistical tests are summarized in table S3.

fig. 1

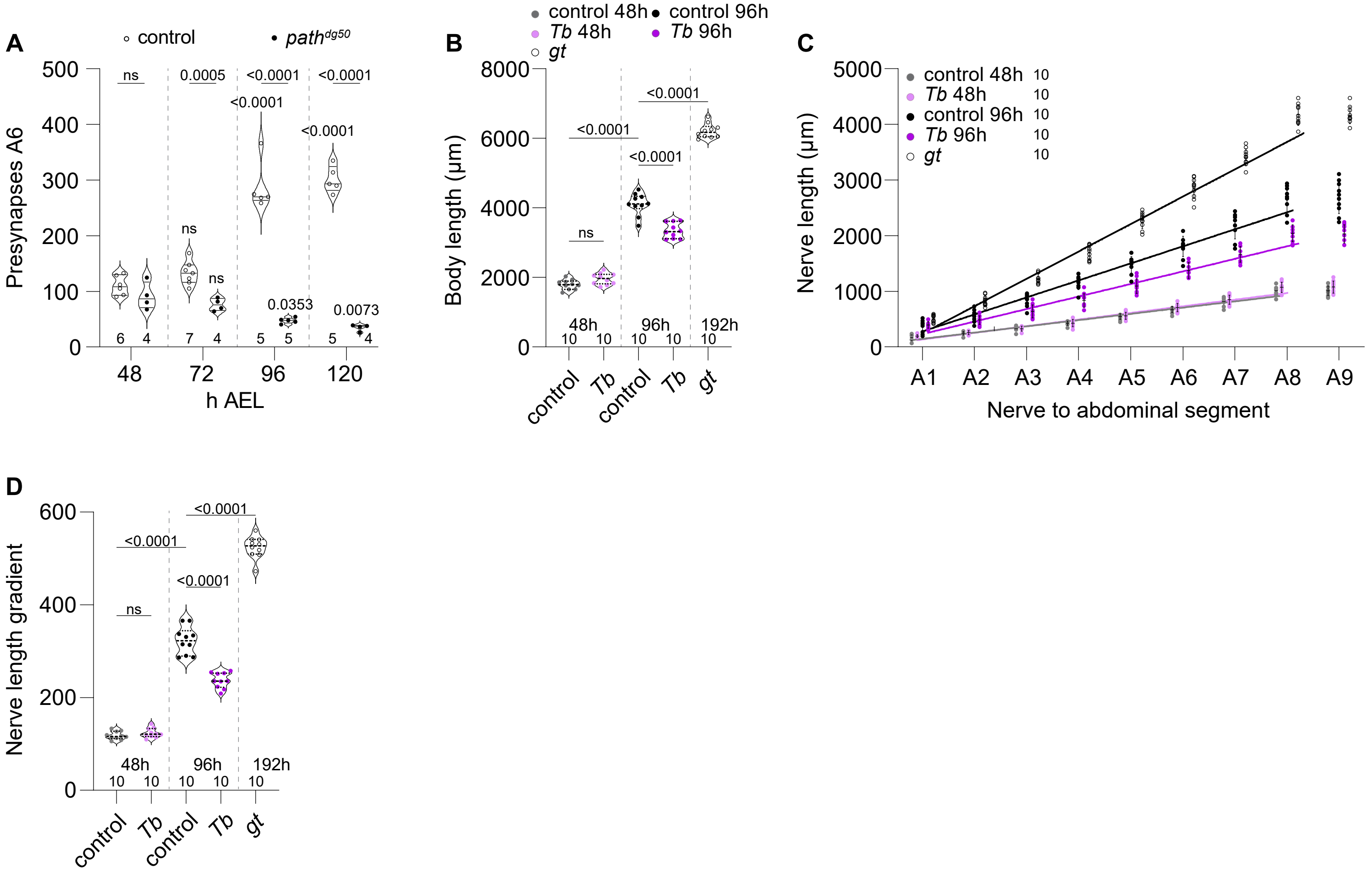

A
control
path^dg50
Presynapses A6
h AEL
ns
0.0005
<0.0001
<0.0001
<0.0001
<0.0001
ns
ns
0.0353
0.0073
B
control 48h
control 96h
Tb 48h
Tb 96h
gt
Body length (µm)
ns
<0.0001
<0.0001
<0.0001
48h
96h
192h
control
Tb
control
Tb
gt
C
control 48h 10
Tb 48h 10
control 96h 10
Tb 96h 10
gt 10
Nerve length (µm)
Nerve to abdominal segment
A1
A2
A3
A4
A5
A6
A7
A8
A9
D
Nerve length gradient
ns
<0.0001
<0.0001
<0.0001
48h
96h
192h
control
Tb
control
Tb
gt

**Supplementary figure 1. Mutation of *path* induces axon length-dependent degeneration.**

**(A)** *path* mutation leads to a progressive loss of presynapses during larval development. Plot shows presynapses from segment A6 from 48 h to 120 h AEL in control and $path^{dg50}$ mutant larvae. *path* mutants exhibit significantly fewer presynapses at 96 h and 120 h AEL in comparison to 48 h AEL, while control larvae exhibit a significant increase in presynapses during development. P-Values, two-way ANOVA with post-hoc Tukey's multiple comparison test. **(B)** *Tb* mutant and *gt* mutant larvae display respectively smaller and larger body length at L3. Body length of control, *Tb* mutant second instar (48 h AEL) and third instar (96h AEL) larvae, and *gt* mutant persistent third instar larvae (192 h AEL). At L2 there is no significant difference in body length between *Tb* mutant and control larvae. During development from L2 to L3 control larvae grow about 2-fold in length, whereas *Tb* mutants grow significantly shorter. *gt* mutants reach 3-fold body length due to prolonged L3 stage. P-Values, ANOVA with post-hoc Šídák's multiple comparison test. **(C-D)** Nerve lengths of animals shown in **(B)**. **(C)** Plot depicts nerve lengths of each abdominal segment (A1-A9) with A8 and A9 sharing the same nerve. Linear regression lines indicate continuously longer nerves for each following segment. **(D)** Nerve length gradients from animals shown in B and C. Inclining slopes from linear regression display significantly larger linear growth demand in posterior neurons than anterior counterparts during development from 48 h-96 h AEL. Axon length correlates with increasing body size: *Tb* mutants display significantly decreased inclining slopes and thereby less growth demand during the same timeframe. Growth demand increases even further in *gt* mutants.  P-Values, ANOVA with post-hoc Šídák's multiple comparison test. Sample numbers are shown for each specimen. Detailed experimental genotypes are provided in table S4.

fig. 2

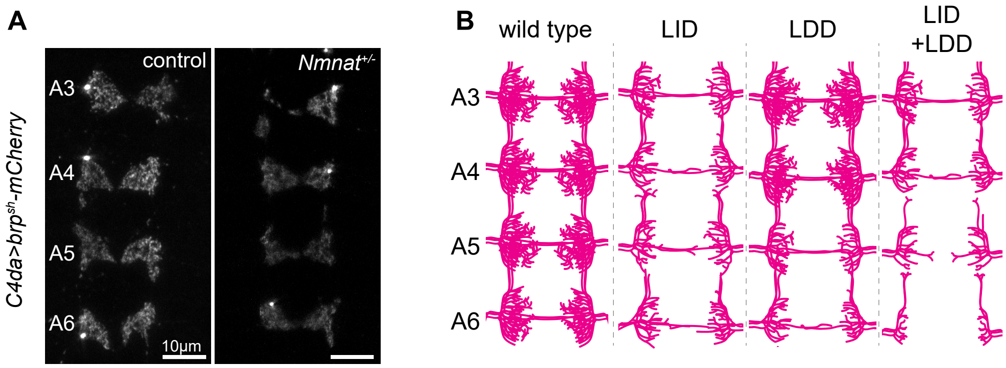

A
control
$Nmnat^{+/-}$
A3
A4
A5
A6
10µm
$C4da>brp^{sh}$-mCherry
B
wild type
LID
LDD
LID
+LDD
A3
A4
A5
A6

**Supplementary figure 2. *Nmnat*$^{+/-}$ mutation induces axon length-independent degeneration.**

**(A)** *Nmnat*$^{+/-}$ mutation leads to decreased size of presynaptic domains that is not affected by axon length. Maximum intensity projections show the distribution of C4da neuron presynapses labeled by *LexAop-brp*$^{sh}$*-mCherry* in the VNC of representative control and *Nmnat*$^{+/-}$ heterozygous larvae. **(B)** Schematic depicting C4da neuron terminal axon domains in segments A3-A6 under the following conditions: wild type, LID, LDD, and cooccurrence LID and LDD. Detailed experimental genotypes are provided in table S4.

fig. 3

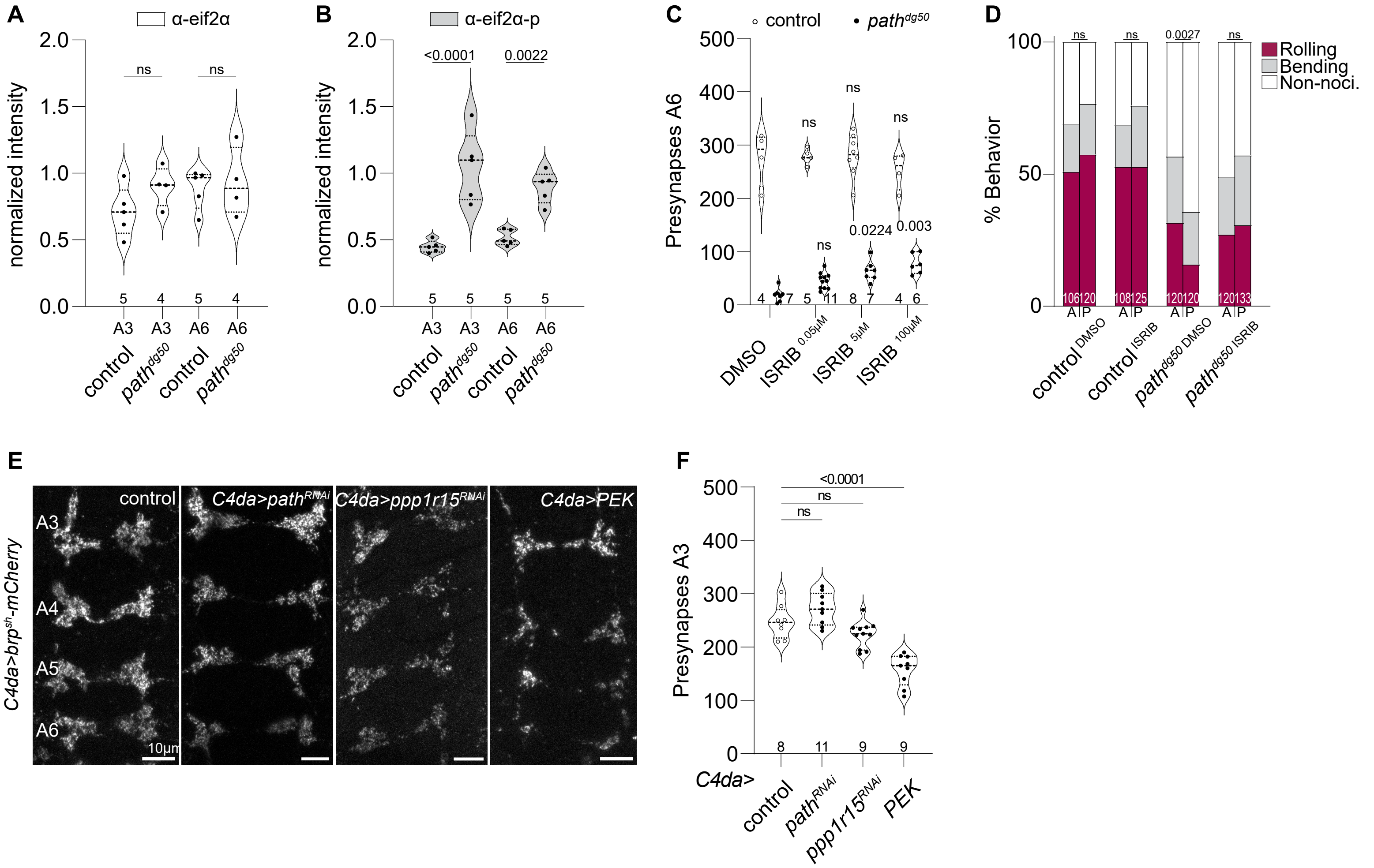
A
α-eif2α
normalized intensity
ns
ns
A3
A3
A6
A6
control
path^dg50
control
path^dg50
B
α-eif2α-p
normalized intensity
<0.0001
0.0022
C
control
path^dg50
Presynapses A6
DMSO
ISRIB 0.05µM
ISRIB 5µM
ISRIB 100µM
0.0224
0.003
D
Rolling
Bending
Non-noci.
% Behavior
0.0027
control DMSO
control ISRIB
path^dg50 DMSO
path^dg50 ISRIB
E
C4da>brp^sh-mCherry
control
C4da>path^RNAi
C4da>ppp1r15^RNAi
C4da>PEK
A3
A4
A5
A6
10µm
F
Presynapses A3
C4da>
control
path^RNAi
ppp1r15^RNAi
PEK
<0.0001

**Supplementary figure 3. Activation of the integrated stress response triggers LDD.** **(A-B)** ISR activation in C4da neurons of *path* mutants. Plots depict normalized immunoreactivity to **(A)** eif2α and **(B)** phosphorylated eif2α (eif2α-P) in C4da neuron somas from segments A3 and A6 of larvae of the indicated genotypes. *path*$^{dg50}$ mutants exhibit significant increases in eif2α-P but not eif2α immunoreactivity. P values, ANOVA with post-hoc Šídák's multiple comparison test. **(C)** The small molecule ISR inhibitor ISRIB inhibits LDD induced by *path* mutation. The plot depicts A6 presynapse numbers of *path*$^{dg50}$ and control larvae fed with the indicated ISRIB concentrations (0.05 μM, 5 μM, 100 μM). ISRIB feeding significantly increased presynapse numbers of *path* mutants when compared to DMSO vehicle controls. P-Values, Two-way ANOVA with post-hoc Tukey's multiple comparison test. **(D)** Axon length-dependent but not length-independent deficits in mechanonociception of *path* mutants are suppressed by ISRIB feeding. Plot depicts behavioral responses of larvae of the indicated genotypes to anterior and posterior noxious mechanical stimuli. P values, Chi-square test. **(E)** Activating the ISR in C4da neurons triggers degeneration in presynaptic domains. Maximum intensity projections depict the distribution of C4da neuron presynapses labeled by *ppk-GAL4>UAS-brp*$^{sh}$*-mCherry* in VNC segments A3-A6 of control larvae, larvae with C4da-specific *path*$^{RNAi}$, or larvae with C4da-specific ISR activation (via *ppp1r15* $^{RNAi}$ or *PEK* overexpression). **(F)** *PEK* overexpression triggers LID. Violin plot depicts A3 presynapse numbers from control larvae, larvae with C4da-specific *path*$^{RNAi}$, or larvae with C4da-specific ISR activation (via *ppp1r15* $^{RNAi}$ or *PEK* overexpression). *PEK* overexpression in C4da neurons leads to significantly lower presynapses numbers. Sample numbers are shown for each specimen. Detailed experimental genotypes are provided in table S4.

fig. 4

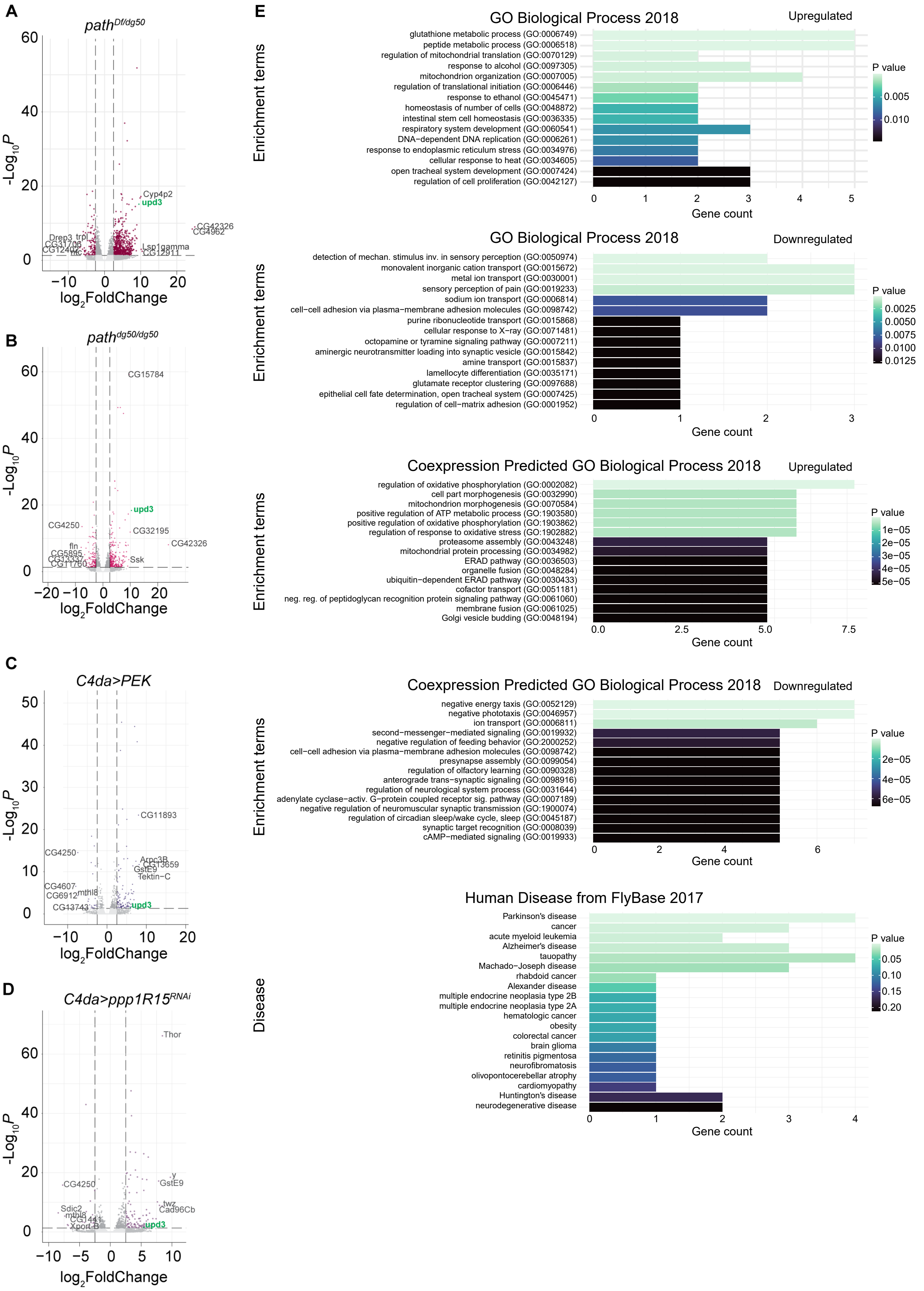
A
path^Df/dg50
-Log10P
log2FoldChange
Cyp4p2
upd3
CG42326
CG4962
Drep3
trpl
CG31706
CG12402
Lsp1gamma
CG12911
B
path^dg50/dg50
CG15784
upd3
CG4250
CG32195
CG42326
fln
CG5895
Ssk
CG11760
C
C4da>PEK
CG11893
CG4250
Arpc3B
CG13659
GstE9
Tektin-C
CG4607
CG6912
mthl8
CG13743
upd3
D
C4da>ppp1R15^RNAi
Thor
CG4250
y
GstE9
Sdic2
mthl8
CG1441
Xport-B
twz
Cad96Cb
upd3
E
GO Biological Process 2018
Upregulated
Enrichment terms
glutathione metabolic process (GO:0006749)
peptide metabolic process (GO:0006518)
regulation of mitochondrial translation (GO:0070129)
response to alcohol (GO:0097305)
mitochondrion organization (GO:0007005)
regulation of translational initiation (GO:0006446)
response to ethanol (GO:0045471)
homeostasis of number of cells (GO:0048872)
intestinal stem cell homeostasis (GO:0036335)
respiratory system development (GO:0060541)
DNA−dependent DNA replication (GO:0006261)
response to endoplasmic reticulum stress (GO:0034976)
cellular response to heat (GO:0034605)
open tracheal system development (GO:0007424)
regulation of cell proliferation (GO:0042127)
Gene count
P value
0.005
0.010
GO Biological Process 2018
Downregulated
detection of mechan. stimulus inv. in sensory perception (GO:0050974)
monovalent inorganic cation transport (GO:0015672)
metal ion transport (GO:0030001)
sensory perception of pain (GO:0019233)
sodium ion transport (GO:0006814)
cell−cell adhesion via plasma−membrane adhesion molecules (GO:0098742)
purine ribonucleotide transport (GO:0015868)
cellular response to X−ray (GO:0071481)
octopamine or tyramine signaling pathway (GO:0007211)
aminergic neurotransmitter loading into synaptic vesicle (GO:0015842)
amine transport (GO:0015837)
lamellocyte differentiation (GO:0035171)
glutamate receptor clustering (GO:0097688)
epithelial cell fate determination, open tracheal system (GO:0007425)
regulation of cell−matrix adhesion (GO:0001952)
Gene count
P value
0.0025
0.0050
0.0075
0.0100
0.0125
Coexpression Predicted GO Biological Process 2018
Upregulated
regulation of oxidative phosphorylation (GO:0002082)
cell part morphogenesis (GO:0032990)
mitochondrion morphogenesis (GO:0070584)
positive regulation of ATP metabolic process (GO:1903580)
positive regulation of oxidative phosphorylation (GO:1903862)
regulation of response to oxidative stress (GO:1902882)
proteasome assembly (GO:0043248)
mitochondrial protein processing (GO:0034982)
ERAD pathway (GO:0036503)
organelle fusion (GO:0048284)
ubiquitin−dependent ERAD pathway (GO:0030433)
cofactor transport (GO:0051181)
neg. reg. of peptidoglycan recognition protein signaling pathway (GO:0061060)
membrane fusion (GO:0061025)
Golgi vesicle budding (GO:0048194)
Gene count
P value
1e−05
2e−05
3e−05
4e−05
5e−05
Coexpression Predicted GO Biological Process 2018
Downregulated
negative energy taxis (GO:0052129)
negative phototaxis (GO:0046957)
ion transport (GO:0006811)
second−messenger−mediated signaling (GO:0019932)
negative regulation of feeding behavior (GO:2000252)
cell−cell adhesion via plasma−membrane adhesion molecules (GO:0098742)
presynapse assembly (GO:0099054)
regulation of olfactory learning (GO:0090328)
anterograde trans−synaptic signaling (GO:0098916)
regulation of neurological system process (GO:0031644)
adenylate cyclase−activ. G−protein coupled receptor sig. pathway (GO:0007189)
negative regulation of neuromuscular synaptic transmission (GO:1900074)
regulation of circadian sleep/wake cycle, sleep (GO:0045187)
synaptic target recognition (GO:0008039)
cAMP−mediated signaling (GO:0019933)
Gene count
P value
2e−05
4e−05
6e−05
Human Disease from FlyBase 2017
Disease
Parkinson's disease
cancer
acute myeloid leukemia
Alzheimer's disease
tauopathy
Machado−Joseph disease
rhabdoid cancer
Alexander disease
multiple endocrine neoplasia type 2B
multiple endocrine neoplasia type 2A
hematologic cancer
obesity
colorectal cancer
brain glioma
retinitis pigmentosa
neurofibromatosis
olivopontocerebellar atrophy
cardiomyopathy
Huntington's disease
neurodegenerative disease
Gene count
P value
0.05
0.10
0.15
0.20

**Supplementary figure 4. RNA-seq analysis of C4da neurons undergoing LDD.**

**(A)** Volcano plot of differentially expressed genes in homozygous *path*$^{dg50}$ mutants compared to heterozygous control. Top 10 DEG indicated in plot with *upd3* (green). padj cutoff = 10e-5, and lfc cutoff = 2.5. total = 13969 variables. **(B)** Volcano plot of differentially expressed genes in *path*$^{dg50}$*/Df* mutants compared to heterozygous control. Top 10 DEG indicated in plot with *upd3* (green). padj cutoff = 10e-5, and lfc cutoff = 2.5. total = 13969 variables. **(C)** Volcano plot of differentially expressed genes in larvae overexpressing *PEK* in C4da neurons compared to control. Top 10 DEG indicated in plot with *upd3* (green). padj cutoff = 10e-5, and lfc cutoff = 2.5. total = 13969 variables. **(D)** Volcano plot of differentially expressed genes in larvae expressing *ppp1R15*$^{RNAi}$ in C4da neurons compared to control. Top 10 DEG indicated in plot with *upd3* (green). padj cutoff = 10e-5, and lfc cutoff = 2.5. total = 13969 variables. **(E)** Gene ontology analysis of the common 109 DEGs indicate deregulation of stress pathways. Plots show gene counts of up- and downregulated genes matching gene ontology terms from Biological Process 2018 and Coexpression Predicted GO Biological Process 2018 as well as Human Disease from Flybase 2017 of all 109 DEGs. Glutathione metabolic process, regulation of mitochondrial translation, response to alcohol, regulation of translation initiation, response to endoplasmic reticulum stress, cellular response to heat, regulation of oxidative phosphorylation, regulation of response to oxidative stess, and organelle fusion indicate active genes involved in cellular stress. Diseases linked to DEGs indicate different Neurodegenerative Diseases which display specific vulnerability of large neurons. Sample numbers are shown for each specimen. Detailed experimental genotypes are provided in table S4.

fig. 5

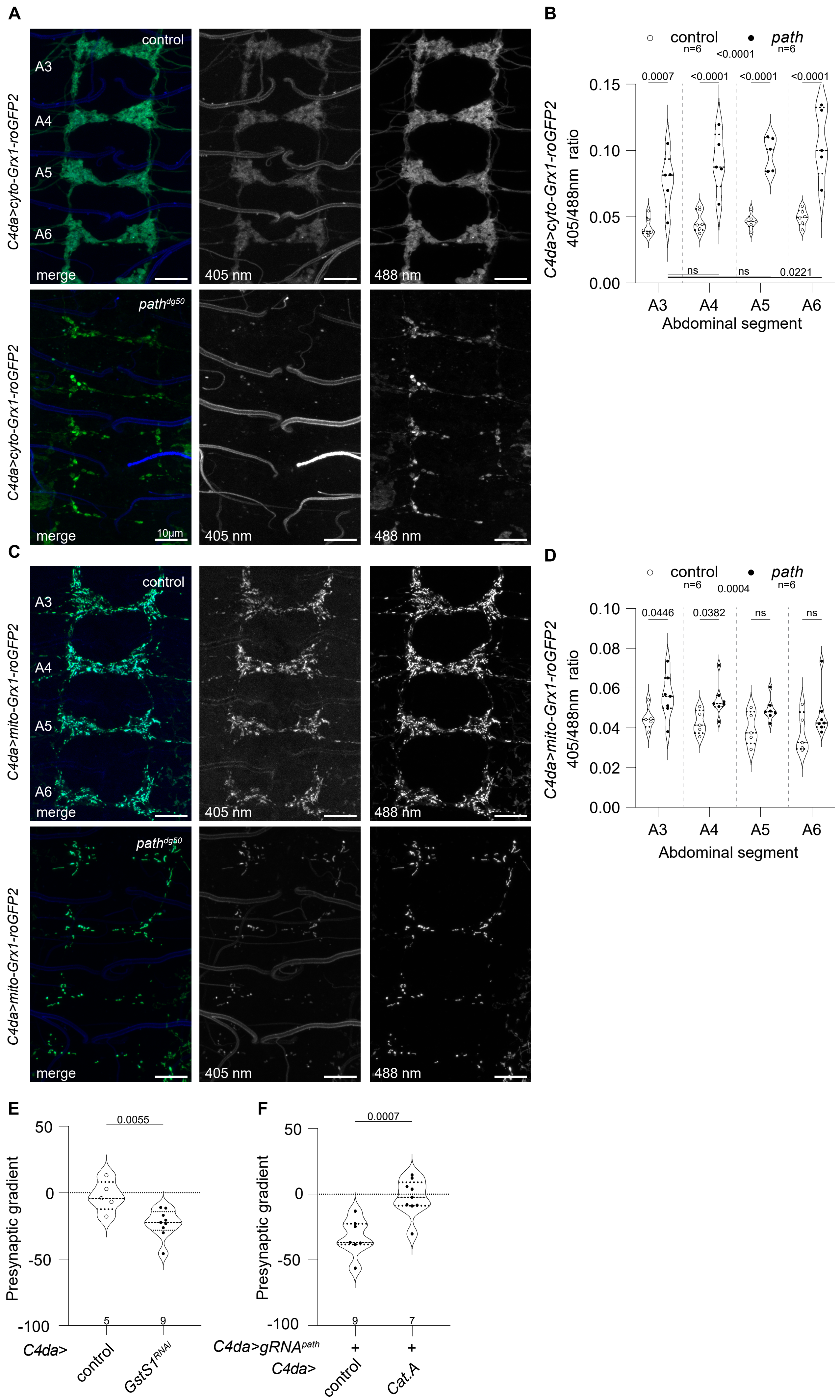
A
control
A3
A4
A5
A6
merge
405 nm
488 nm
C4da>cyto-Grx1-roGFP2
path^dg50
10µm
B
control n=6
path n=6
<0.0001
0.0007
<0.0001
<0.0001
<0.0001
ns
ns
0.0221
C4da>cyto-Grx1-roGFP2 405/488nm ratio
Abdominal segment
C
C4da>mito-Grx1-roGFP2
D
0.0004
0.0446
0.0382
ns
ns
C4da>mito-Grx1-roGFP2 405/488nm ratio
E
0.0055
Presynaptic gradient
5
9
C4da>
control
GstS1^RNAi
F
0.0007
9
7
C4da>gRNA^path
+
+
C4da>
control
Cat.A

**Supplementary figure 5.** ***Defects in redox potential contribute to LDD.***

**(A-D)** Glutathione redox potential is decreased in *path* mutants. **(A)** Maximum intensity projections depict A3-A6 presynaptic domains labeled by expression of the ratiometric cytosolic redox reporter *UAS-cyto-Grx1-roGFP2* in C4da neurons*.* Images depict signal from excitation with 405 or 488 nm laser lines, as indicated, or a merge of the two channels in *path* mutant and wild-type control larvae. **(B)** Violin plots depict cyto-Grx1-roGFP2 signal intensity ratios (405/488 nm excitation) from the presynaptic domains in segments A3-A6 of control and *path* mutant larvae. $path^{dg50}$ mutants exhibit a strong increase in cytosolic 405/488 nm ratios in all segments compared to wild-type controls, indicative of a decrease potential of cells to fight oxidative stress in C4da neurons. P-Values, Two-way ANOVA with post-hoc Tukey's multiple comparison test. **(C)** Maximum intensity projections depict A3-A6 presynaptic domains labeled by expression of the ratiometric mitochondrial redox reporter *UAS-mito-Grx1-roGFP2* in C4da neurons of *path* mutant and wild-type control larvae*.* **(D)** Violin plots depict mito-Grx1-roGFP2 405/488 nm intensity ratios in VNC segments A3-A6. $path^{dg50}$ displays robust increases in mitochondrial 405/488 nm ratios in comparison to wild-type controls, indicative of a decreased potential of mitochondria to fight oxidative stress in C4da neurons. P-Values, Two-way ANOVA with post-hoc Tukey's multiple comparison test. **(E)** Knockdown of *GstS1* in C4da neurons leads to LDD. Violin plot shows presynaptic gradient measurements of control larvae and larvae expressing $GstS1^{RNAi}$ in C4da neurons. *GstS1* knockdown induces a significant change in the presynaptic gradient. P-value, unpaired t-test. **(F)** *Catalase* overexpression rescues LDD. Violin plot shows effects of *Catalase* (*Cat.A*) expression on presynaptic gradient triggered by CRISPR/Cas9-mediated *path* inactivated in C4da neurons *Cat.A* overexpression significantly attenuates the presynaptic gradient triggered by *path* inactivation. P-value, unpaired t-test. Sample numbers are shown for each specimen. Detailed experimental genotypes are provided in table S4.

fig. 6

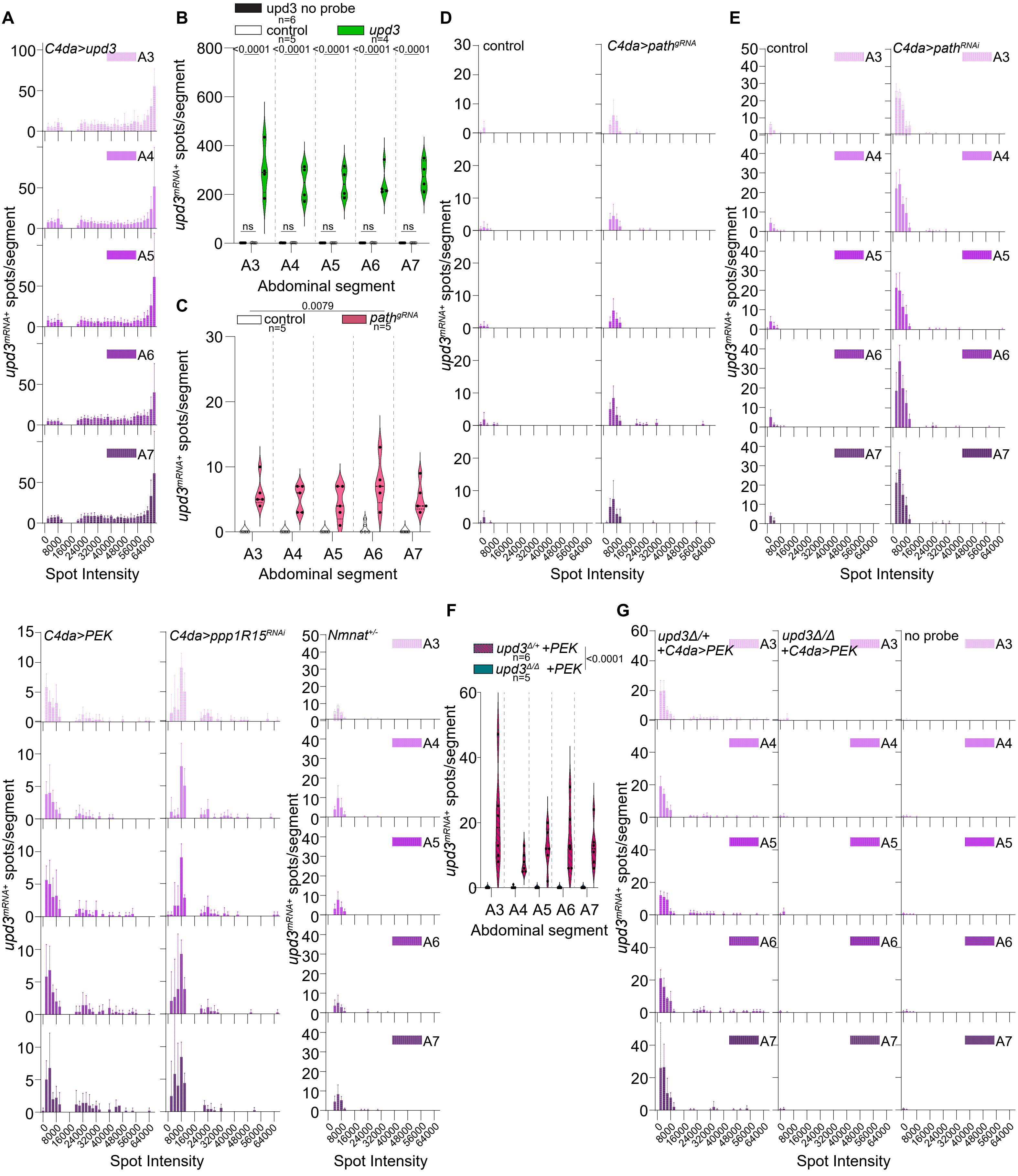

A
B
C
D
E
F
G
C4da>upd3
upd3 no probe
control
upd3
path^gRNA
C4da>path^gRNA
C4da>path^RNAi
C4da>PEK
C4da>ppp1R15^RNAi
Nmnat^+/-
upd3^Δ/+ +PEK
upd3^Δ/Δ +PEK
upd3Δ/+ +C4da>PEK
upd3Δ/Δ +C4da>PEK
no probe
A3
A4
A5
A6
A7
Abdominal segment
Spot Intensity
upd3^mRNA+ spots/segment
<0.0001
0.0079
ns
n=6
n=5
n=4

**Supplementary figure 6.** ***HCR-FISH analysis of upd3 expression in C4da neurons.***
**(A-B)** *upd3$^{mRNA}$* probes targeting *upd3*: *upd3* overexpression in C4da leads to strong detection of *upd3* puncta. **(A)** Histogram depicts the number and intensity of *upd3* HCR-FISH puncta in C4da neuron soma from the indicated abdominal segments in larvae overexpressing *upd3* in C4da neurons. **(B)** Violin plots depict the number of *upd3* puncta (intensity>8000) in each abdominal segment of neurons overexpressing *upd3. upd3* overexpression in C4da leads to a comparable increase of *upd3* puncta in all segments. P-Values, Two-way ANOVA with post-hoc Dunnett's multiple comparison test. **(C-D)** *path* knockdown cell-autonomously increases *upd3* expression in C4da neurons. **(C)** Violin plots depict the number of *upd3* puncta (intensity>8000) in C4da neurons from the indicated abdominal segments of control or *path* knockdown (*ppk-CAS9, path$^{gRNA}$*) larvae. Loss of *path* in C4da neurons leads to a significant increase of *upd3* puncta compared to controls. P-value, Mann Whitney test. **(D)** Histograms depict the number and intensity of *upd3* HCR-FISH puncta in C4da neuron soma of specimens depicted in **(C)**. **(E)** Treatments that induce the ISR in C4da neurons trigger *upd3* expression (companion to Fig. 2, D and E). Histograms depict the number and intensity of *upd3* HCR-FISH puncta in C4da neuron soma from the abdominal segments of the indicated specimens. **(F-G)** Specificity of *upd3$^{mRNA}$* probes. **(F)** Violin plots depict the number of *upd3* puncta (intensity>8000) in C4da neurons from the indicated segments in larvae overexpressing *PEK* in C4da neurons and additionally heterozygous or homozygous for a *upd3Δ* mutation, as indicated. *PEK* overexpression in C4da leads to significant increase of *upd3* puncta in all segments within a heterozygous *upd3Δ* mutant background. In a homozygous background *PEK* overexpression no *upd3* puncta can be detected. Similarly, no puncta are detectable specimens stained without a *upd3* probe. P-values, Kruskal-Wallis test with post-hoc Dunn's test. **(G)** Histograms depict the number and intensity of *upd3* HCR-FISH puncta in C4da neuron soma from the indicated abdominal segments in specimens indicated in **(F)**. **(H)** Histograms depict the number and intensity of *upd3* HCR-FISH puncta in C4da neuron soma from the indicated segments of *Nmnat$^{+/-}$* heterozygous mutant larvae. Sample numbers are shown for each specimen. Detailed experimental genotypes are provided in table S4.

fig. 7

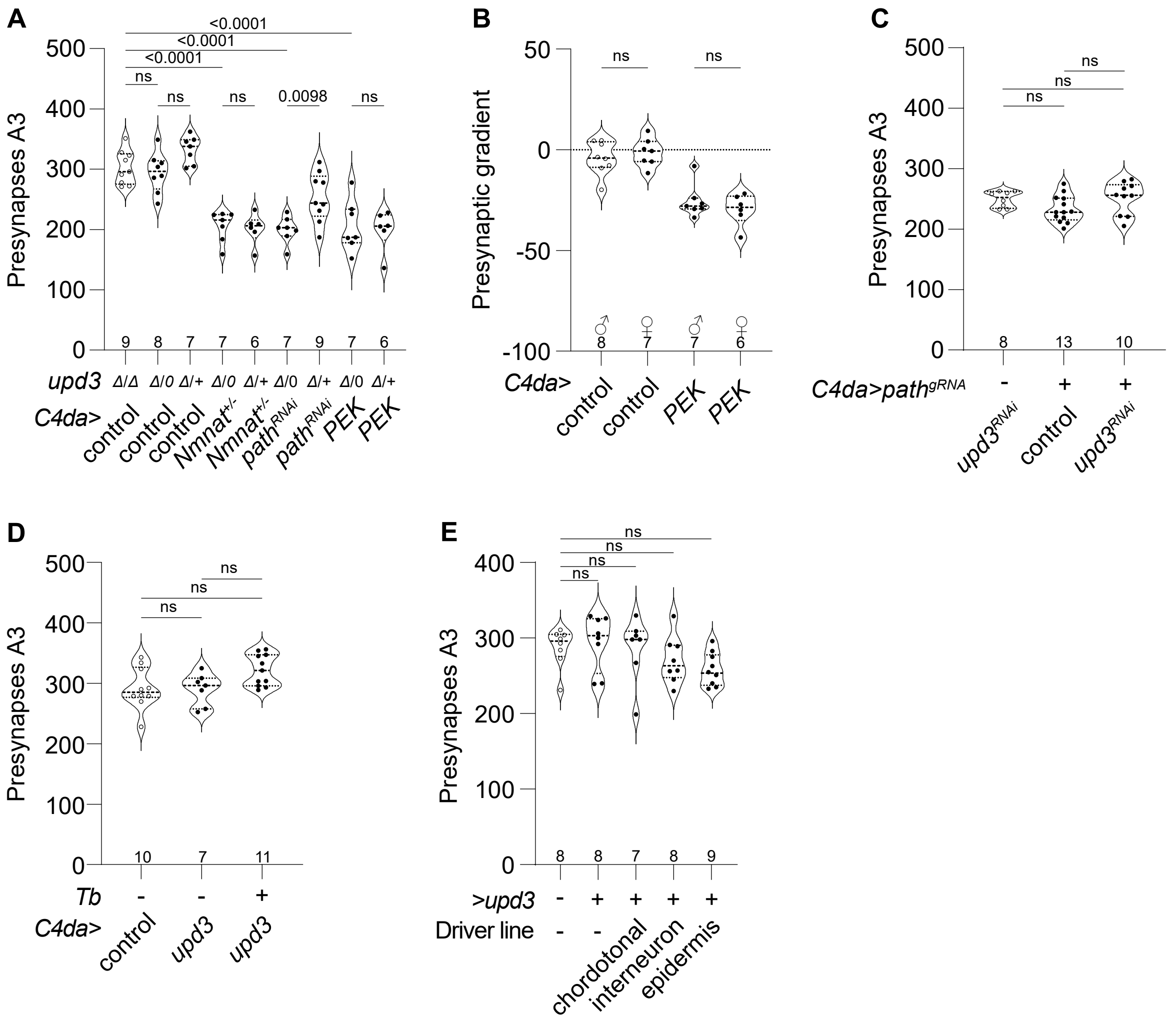
A
Presynapses A3
<0.0001
ns
0.0098
upd3
C4da>
control
Nmnat+/-
pathRNAi
PEK
B
Presynaptic gradient
C4da>
control
PEK
C
C4da>pathgRNA
upd3RNAi
control
D
Tb
C4da>
upd3
E
>upd3
Driver line
chordotonal
interneuron
epidermis

**Supplementary figure 7. *upd3* is dispensable for LID.**

**(A)** Mutation of upd3 has no effect on presynapse loss phenotypes in segment A3. Violin plots depict A3 presynapse numbers of genotypes indicated in Fig. 2F. The LID phenotypes triggered by *Nmnat* mutation, $path^{RNAi}$ or *PEK* overexpression are unaffected by loss of *upd3.* P-Values, ANOVA with post-hoc Šídák's multiple comparison test. **(B)** Violin plots depict presynapse gradient measurements from control larvae and larvae overexpressing *PEK* in C4da neurons. Comparison of males and females revealed no significant difference in C4da presynapse numbers or slope. P-values, Kruskal-Wallis test with post-hoc Dunn’s test. **(C)** Knockdown of *upd3* in C4da neurons has no effect on LID induced by *path* KO in C4da neurons. Violin plots depict A3 presynapse numbers of genotypes indicated in Fig. 2G. P-Values, ANOVA with post-hoc Tukey's multiple comparison test. **(D)** Overexpression of *upd3* in C4da neurons has no effect on A3 presynapse numbers in control or *Tb* mutant larvae. Violin plots depict A3 presynapse numbers of genotypes indicated in Fig. 2I. P-Values, ANOVA with post-hoc Tukey's multiple comparison test. **(E)** Overexpression of *upd3* in cho neurons, ABLK interneurons, or epidermal cells has no effect on A3 presynapse numbers in control or *Tb* mutant larvae. Violin plots depict A3 presynapse numbers of genotypes indicated in Fig. 2L. P-Values, ANOVA with post-hoc Dunnett's multiple comparison test. Sample numbers are shown for each specimen. Detailed experimental genotypes are provided in table S4.

fig. 8

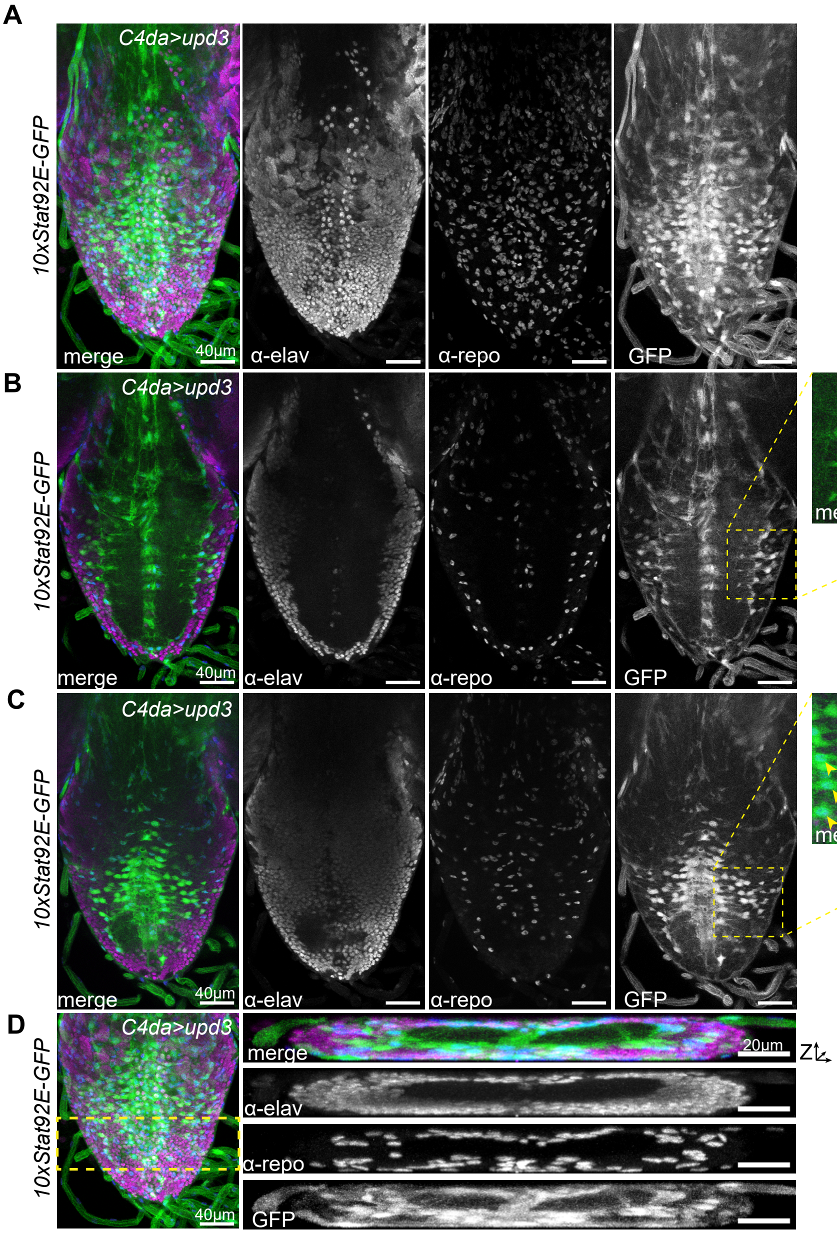
A
B
C
D
C4da>upd3
10xStat92E-GFP
merge
α-elav
α-repo
GFP
40µm
20µm
Z

**Supplementary figure 8. *upd3* expression in C4da neurons induces Stat92E activity in the VNC. (A-D)** Representative images depict *10xStat92E-GFP* reporter activity (GFP immunoreactivity, green) in VNC preparations additionally labeled for repo (glial marker, blue) and elav (neuronal marker, magenta) immunoreactivity. Larvae overexpress *upd3* selectively in C4da neurons. **(A)** Maximum intensity projection of a confocal volume containg the entire VNC. **(B)** Maximum intensity projection of a confocal volume containing upper medial slices through the neuropil. Zoomed view of the ROI (yellow hatched box) shows high resolution view of antibody labeling. White arrows, elav-positive GFP-expressing cells; yellow arrows, repo-positive GFP-expressing cells. **(C)** Maximum intensity projection of a confocal volume containing lower medial slices to the ventral neuropil. **(D)** XZ-projection showing *10xStat92E-GFP* reporter activity in the ventral portion of the VNC. Detailed experimental genotypes are provided in table S4.

fig. 9

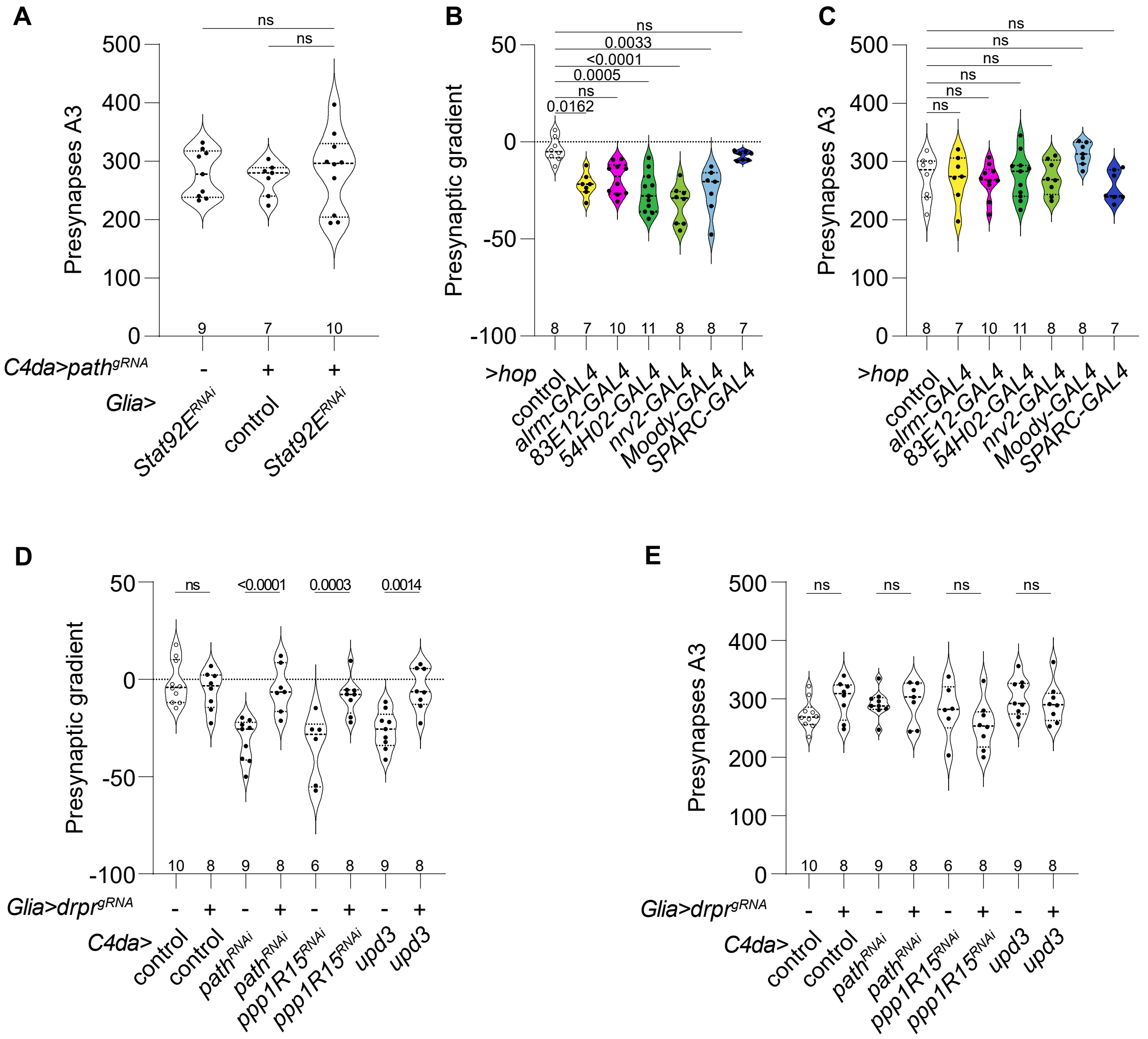
A
Presynapses A3
ns
ns
9
7
10
C4da>path$^{gRNA}$
-
+
+
Glia>
Stat92E$^{RNAi}$
control
Stat92E$^{RNAi}$
B
Presynaptic gradient
ns
0.0033
<0.0001
0.0005
ns
0.0162
8 7 10 11 8 8 7
>hop
control
alrm-GAL4
83E12-GAL4
54H02-GAL4
nrv2-GAL4
Moody-GAL4
SPARC-GAL4
C
Presynapses A3
ns
ns
ns
ns
ns
ns
8 7 10 11 8 8 7
>hop
control
alrm-GAL4
83E12-GAL4
54H02-GAL4
nrv2-GAL4
Moody-GAL4
SPARC-GAL4
D
Presynaptic gradient
ns
<0.0001
0.0003
0.0014
10 8 9 8 6 8 9 8
Glia>drpr$^{gRNA}$
- + - + - + - +
C4da>
control
control
path$^{RNAi}$
path$^{RNAi}$
ppp1R15$^{RNAi}$
ppp1R15$^{RNAi}$
upd3
upd3
E
Presynapses A3
ns
ns
ns
ns
10 8 9 8 6 8 9 8
Glia>drpr$^{gRNA}$
- + - + - + - +
C4da>
control
control
path$^{RNAi}$
path$^{RNAi}$
ppp1R15$^{RNAi}$
ppp1R15$^{RNAi}$
upd3
upd3

**Supplementary figure 9. *Stat92E* pathway components are necessary and sufficient for LDD but not LID.**

**(A)** Glial knockdown of *Stat92E* has no effect on A3 presynapse numbers in control larvae or larvae with *path* inactivated in C4da neurons. Violin plot depicts A3 presynapse numbers of genotypes indicated in Fig. 3E. P-Values, ANOVA with post-hoc Šídák's multiple comparison test. **(B-C)** Glial overexpression of *hop* triggers LDD phenotypes. Violin plots depict **(B)** presynapse gradient measurements and **(C)** A3 presynapse numbers of larvae in which *hop* was expressed with the indicated GAL4 drivers. different glial subset drivers. Overexpression of *hop* with *alrm-GAL4* (AG)*, 54H02-GAL4* (CG), *nrv2-GAL4* (CG) and *moody-GAL4* (SPG) lead to significant LDD phenotypes in C4da presynapses. P-values, Kruskal-Wallis test with post-hoc Dunn's test. **(D-E)** The engulfment receptor *drpr* is required in glial cells for LDD. Violin plots depict **(D)** presynapse gradient measurements and **(E)** A3 presynapse numbers from larvae in which the indicated transgenes were expressed in C4da neurons to induce LDD and $drpr^{gRNA}$ was used to inactivate *drpr* in glial cells. P-Values, ANOVA with post-hoc Šídák's multiple comparison test, comparing samples with and without glial *drpr* inactivation. Sample numbers are shown for each specimen. Detailed experimental genotypes are provided in table S4.

fig. 10

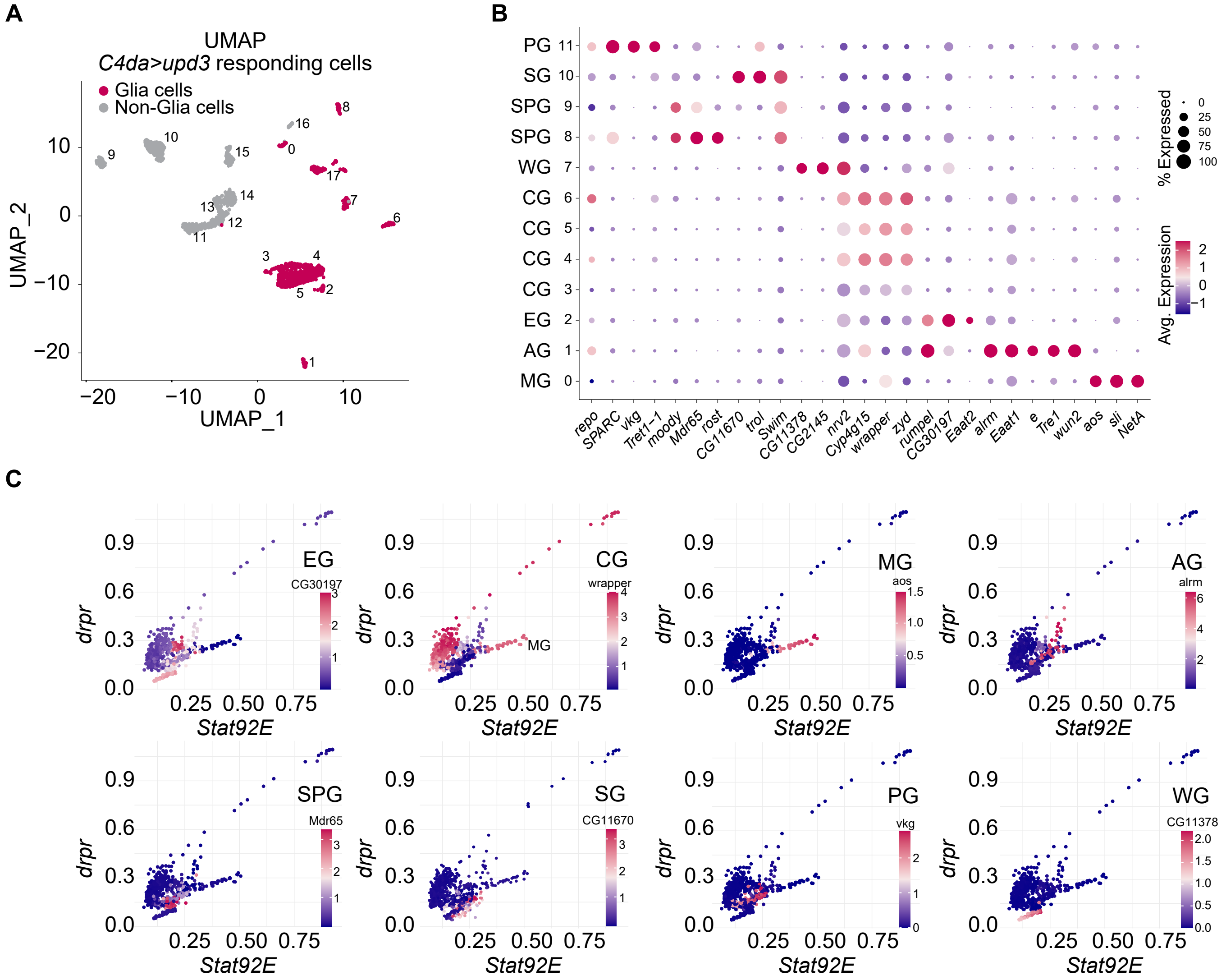
A
UMAP
C4da>upd3 responding cells
Glia cells
Non-Glia cells
UMAP_1
UMAP_2
B
PG 11
SG 10
SPG 9
SPG 8
WG 7
CG 6
CG 5
CG 4
CG 3
EG 2
AG 1
MG 0
repo
SPARC
vkg
Tret1-1
moody
Mdr65
rost
CG11670
trol
Swim
CG11378
CG2145
nrv2
Cyp4g15
wrapper
zyd
rumpel
CG30197
Eaat2
alrm
Eaat1
e
Tre1
wun2
aos
sli
NetA
% Expressed
Avg. Expression
C
EG
CG30197
CG
wrapper
MG
aos
AG
alrm
SPG
Mdr65
SG
CG11670
PG
vkg
WG
CG11378
drpr
Stat92E

**Supplementary figure 10.** ***Upd3 expression in C4da neurons induces high levels of Stat92E expression in a subset of cortex glia***.

**(A)** Initial clustering of 1701 *upd3*-responsive VNC cells yielded 18 cell clusters, including 8 non-glia (grey) and 10 glia clusters (red). **(B)** Glia cluster were subsetted and reclustered, and the dot plot depicts expression of marker genes used to identify the 12 clusters of *upd3*-responsive VNC glia. Dot size indicates the proportion of cells in each cluster expressing a given marker and dot shading indicates mean expression across the cluster. Glia subsets include perineurial glia (PG), surface glia (SG), subperineurial glia (SPG), wrapping glia (WG), cortex glia (CG), ensheathing glia (EG), astrocyte-like glia (AG), and midline glia (MG). **(C)** Expression plots depict expression levels of *drpr* (x-axis) plotted against *Stat92E* (y-axis) for all glial cells. In each plot cells are shaded according to expression levels of the indicated glial subtype marker. CG and MG show highly correlation expression of *drpr* and *Stat92E*, with a subset of CGs displaying strongest expression of both.

fig. 11

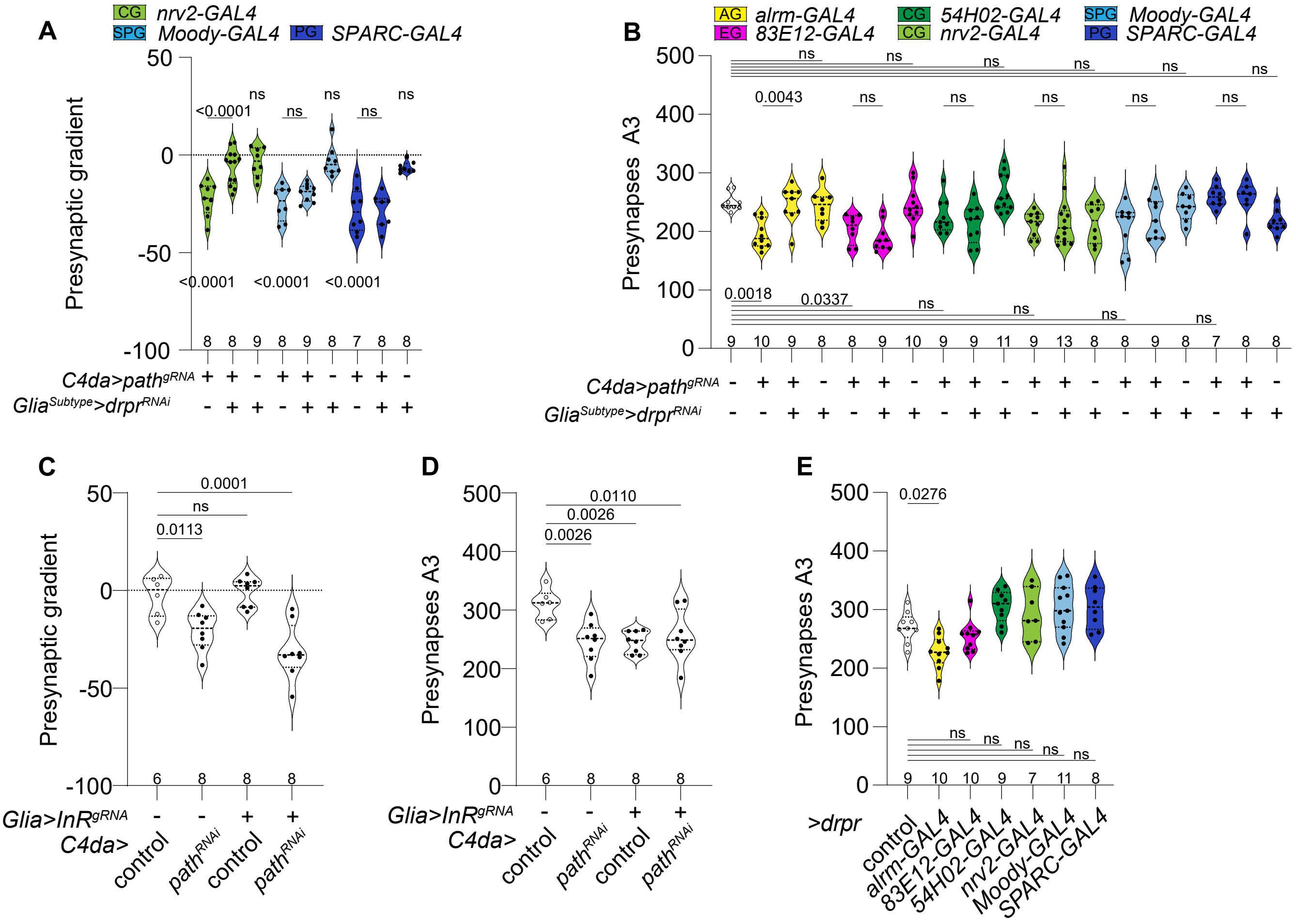

A
CG nrv2-GAL4
SPG Moody-GAL4
PG SPARC-GAL4
Presynaptic gradient
C4da>path^gRNA
Glia^Subtype>drpr^RNAi
B
AG alrm-GAL4
EG 83E12-GAL4
CG 54H02-GAL4
CG nrv2-GAL4
SPG Moody-GAL4
PG SPARC-GAL4
Presynapses A3
C4da>path^gRNA
Glia^Subtype>drpr^RNAi
C
Presynaptic gradient
Glia>InR^gRNA
C4da>
control
path^RNAi
D
Presynapses A3
Glia>InR^gRNA
C4da>
control
path^RNAi
E
Presynapses A3
>drpr
control
alrm-GAL4
83E12-GAL4
54H02-GAL4
nrv2-GAL4
Moody-GAL4
SPARC-GAL4

**Supplementary figure 11. *drpr* functions in cortex glia to mediate LDD.**

**(A)** *drpr* is required in cortex glia for LDD. Violin plot depicts effects of *drpr*$^{RNAi}$ in glial subsets on LDD induced by CRISPR/CAS-9 *path* knockout selectively in C4da neurons. *drpr*$^{RNAi}$ in CG but not AG, EG, SPG, or PG significantly ameliorates presynaptic gradient induced by *path* inactivation (see also Fig. 3L). P-values, Kruskal-Wallis test with post-hoc Dunn's test. **(B)** *drpr* is required in astrocyte-like glia for LID. Violin plot depicts effects of *drpr*$^{RNAi}$ in glial subsets on A3 presynapse numbers in control larvae and larvae in which *path* is selectively knocked out in C4da neurons. *drpr* RNAi in AG but not CG, EG, SPG, or PG significantly ameliorates A3 presynapses induced by *path* inactivation in C4da neurons. P-Values, ANOVA with post-hoc Šídák's multiple comparison test. **(C)** KO of *InR* in glia cells has no effect on LDD induced by *path* KD in C4da neurons. Violin plot depicts presynapse gradient measurements in larvae of the indicated genotypes. P-Values, ANOVA with post-hoc Dunnett's multiple comparison test. **(D)** Loss of *InR* in glia cells reduces presynapse numbers but has no effect on LDD induced by *path* KD in C4da neurons. P-Values, ANOVA with post-hoc Dunnett's multiple comparison test. **(E)** Effects of *drpr* overexpression in different glia subtypes on A3 presynapse numbers. Overexpressing *drpr* with *alarm-GAL4* (AG) but not drivers for other glia subtypes significantly reduced A3 presynapse numbers. P-Values, ANOVA with post-hoc Dunnett's multiple comparison test. Sample numbers are shown for each specimen. Detailed experimental genotypes are provided in table S4.

fig. 12

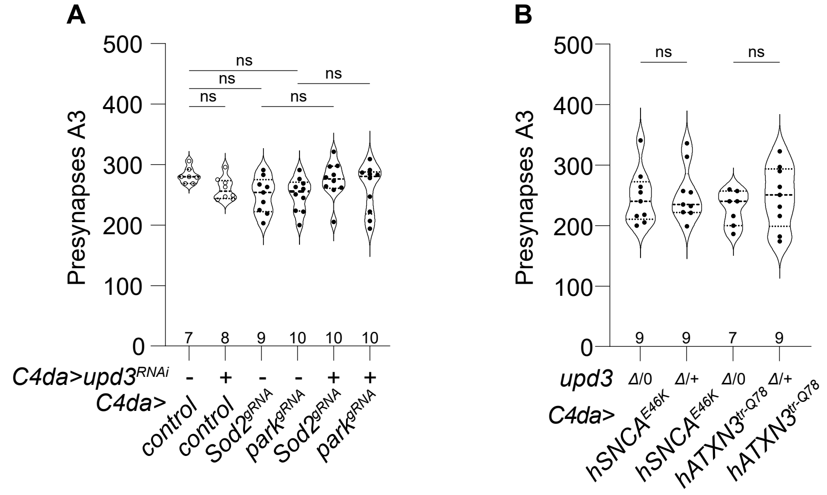
A
Presynapses A3
500
400
300
200
100
0
ns
ns
ns
ns
ns
7 8 9 10 10 10
C4da>upd3RNAi - + - - + +
C4da>
control
control
Sod2gRNA
parkgRNA
Sod2gRNA
parkgRNA
B
Presynapses A3
ns
ns
9 9 7 9
upd3 Δ/0 Δ/+ Δ/0 Δ/+
C4da>
hSNCAE46K
hSNCAE46K
hATXN3tr-Q78
hATXN3tr-Q78

**Supplementary figure 12. Human NDD-associated genetic models do not drive LID in C4da neurons.**

**(A)** Violin plot depicts A3 presynapse numbers of genotypes indicated in Fig. 4 E. No differences in A3 presynapses between *upd3* KD and control with simultanous KO of *Sod2*, or *park.* P-Values, ANOVA with post-hoc Šídák's multiple comparison test. **(B)** Violin plot depicts A3 presynapse numbers of genotypes indicated in Fig. 4 F. No differences in A3 presynapse numbers in *upd3Δ* hemizygous compared to homozygous background during overexpression of $hATXN3^{tr\text{-}Q78}$ *or* $hSNCA^{E46K}$ in C4da neurons. P-Values, ANOVA with post-hoc Šídák's multiple comparison test. Sample numbers are shown for each specimen. Detailed experimental genotypes are provided in table S4.

fig. 13

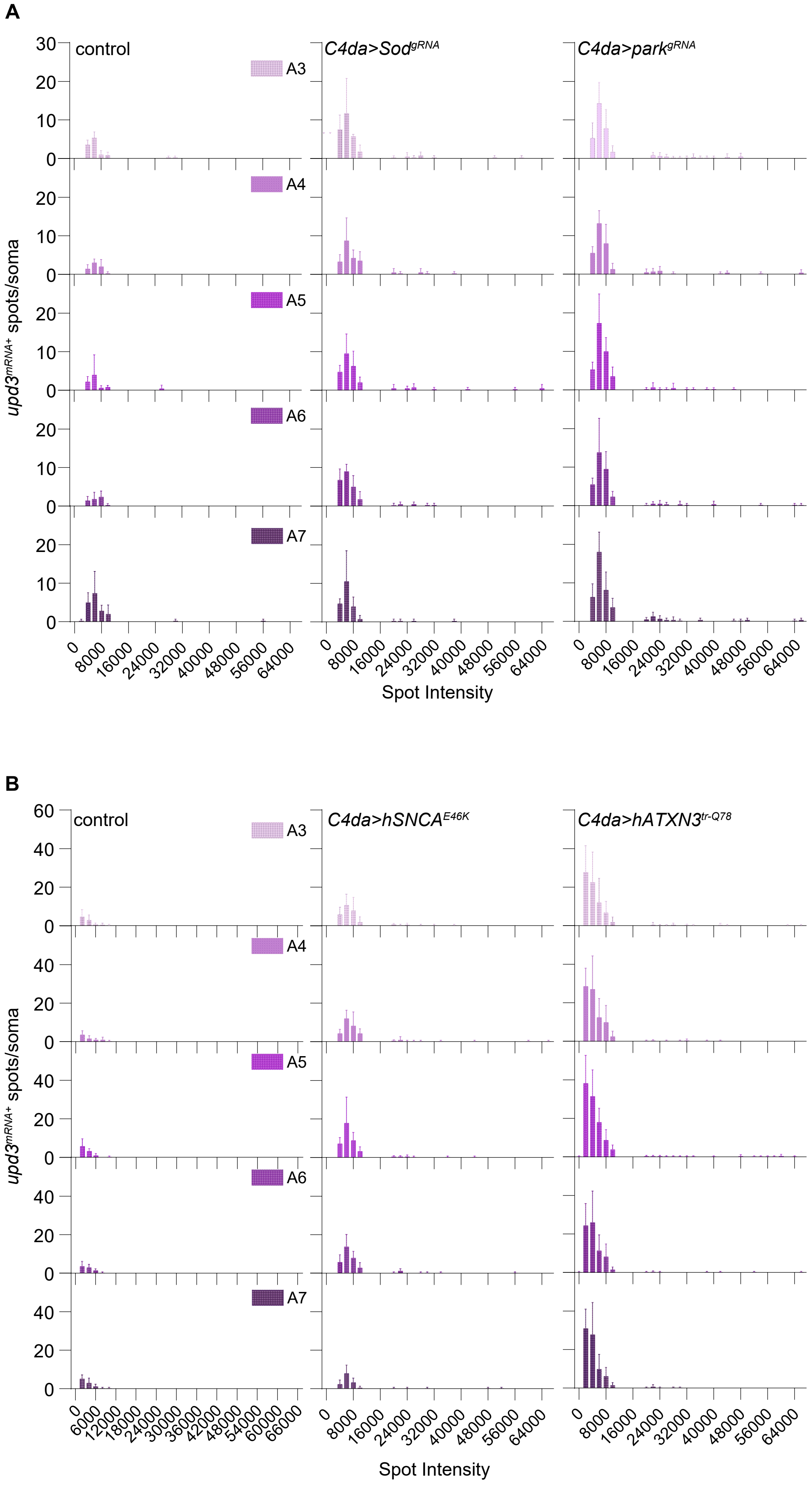
A
control
C4da>Sod$^{gRNA}$
C4da>park$^{gRNA}$
A3
A4
A5
A6
A7
upd3$^{mRNA+}$ spots/soma
0 8000 16000 24000 32000 40000 48000 56000 64000
Spot Intensity
B
control
C4da>hSNCA$^{E46K}$
C4da>hATXN3$^{tr\text{-}Q78}$
A3
A4
A5
A6
A7
upd3$^{mRNA+}$ spots/soma
0 6000 12000 18000 24000 30000 36000 42000 48000 54000 60000 66000
0 8000 16000 24000 32000 40000 48000 56000 64000
Spot Intensity

**Supplementary figure 13. Human NDD-associated genetic models induce *upd3* expression in C4da neurons.**

**(A)** Histograms depict the number and intensity of *upd3* HCR-FISH puncta in C4da neuron soma in segments A3-A7 for each genotype shown in Fig. 4G. **(B)** Histograms depict the number and intensity of *upd3* HCR-FISH puncta in C4da neuron soma in segments A3-A7 for each genotype indicated in Fig. 4H. Detailed experimental genotypes are provided in table S4.

fig. 14

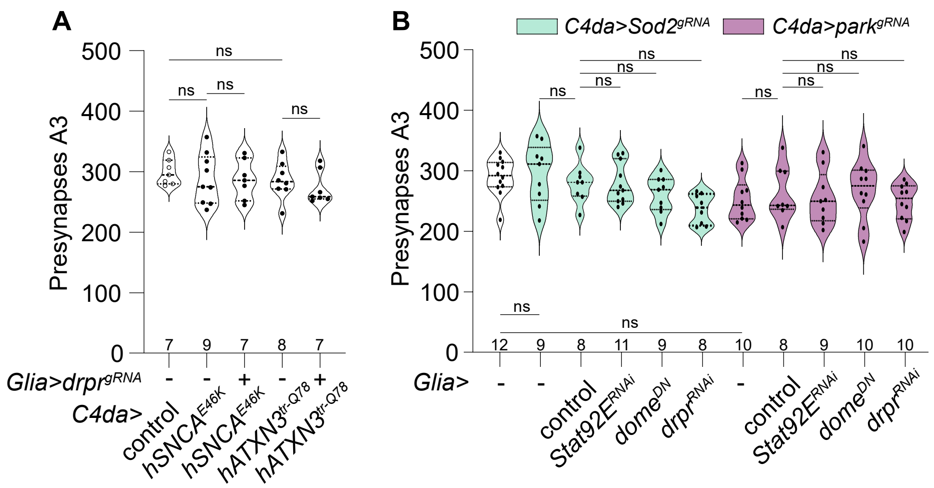
A
B
C4da>Sod2gRNA
C4da>parkgRNA
Presynapses A3
500
400
300
200
100
0
ns
7 9 7 8 7
Glia>drprgRNA
- - + - +
C4da>
control
hSNCAE46K
hSNCAE46K
hATXN3tr-Q78
hATXN3tr-Q78
12 9 8 11 9 8 10 8 9 10 10
Glia>
- -
control
Stat92ERNAi
domeDN
drprRNAi
-
control
Stat92ERNAi
domeDN
drprRNAi

**Supplementary figure 14. Human NDD-associated genetic models do not induce LID in C4da neurons.**

**(A)** Overexpression of *hATXN3*$^{tr\text{-}Q78}$ *or hSNCA*$^{E46K}$ in C4da neurons does not induce LID. Violin plot depicts A3 presynapse numbers for samples depicted in Fig. 4I. P-values, Kruskal-Wallis test with post-hoc Dunn’s test. **(B)** Knockdown of *Stat92E* pathway components does not alter A3 presynapse numbers. Violin plot depicts A3 presynapse numbers of genotypes indicated in Fig. 4J. P-Values, ANOVA with post-hoc Šídák's multiple comparison test. Sample numbers are shown for each specimen. Detailed experimental genotypes are provided in table S4.
